# An ortholog of *P. falciparum* chloroquine resistance transporter (PfCRT) plays a key role in maintaining the integrity of the endolysosomal system in *Toxoplasma gondii* to facilitate host invasion

**DOI:** 10.1101/409904

**Authors:** L. Brock Thornton, Paige Teehan, Katherine Floyd, Christian Cochrane, Amy Bergmann, Bryce Riegel, Paul D. Roepe, Zhicheng Dou

## Abstract

*Toxoplasma gondii* is an apicomplexan parasite with the ability to use foodborne, zoonotic, and congenital routes of transmission that causes severe disease in immunocompromised patients. The parasites harbor a lysosome-like digestive vacuole, termed the “Vacuolar Compartment/Plant-Like Vacuole” (VAC/PLV), which plays an important role in maintaining the lytic cycle and virulence of *T. gondii*. The VAC supplies proteolytic enzymes that are required to mature the parasite’s invasion effectors and that digest autophagosomes and endocytosed host proteins. Previous work identified a *T. gondii* ortholog of the *Plasmodium falciparum* chloroquine resistance transporter (PfCRT) that localized to the VAC. Here, we show that TgCRT is a membrane transporter that is functionally similar to PfCRT. We also genetically ablate *TgCRT* and reveal that TgCRT protein plays a key role in maintaining the integrity of the parasite’s endolysosomal system by controlling morphology of the VAC. When TgCRT is absent, the VAC dramatically increases in size by ~15-fold and co-localizes with its adjacent endosome-like compartment. Presumably to reduce aberrant swelling, transcription and translation of endolysosomal proteases are decreased in Δ*TgCRT* parasites. Expression of one endolysosomal subtilisin protease is quite significantly reduced, which impedes trimming of micronemal proteins, and significantly decreases parasite invasion. Chemical and genetic inhibition of proteolysis within the VAC reverses these effects, reducing VAC size and partially restoring the endolysosomal system, micronemal protein trimming, and invasion. Taken together, these findings reveal for the first time a physiological role of TgCRT in controlling VAC volume and the integrity of the endolysosomal system in *T. gondii*.

**Author Summary:** *Toxoplasma gondii* is an obligate intracellular protozoan parasite that belongs to the phylum Apicomplexa and that infects virtually all warm-blooded organisms. Approximately one-third of the human population is infected with *Toxoplasma*. The parasites invade host cells via processed invasion effectors in order to disseminate infection. A lysosome-like digestive vacuole (VAC) is involved in refining these invasion effectors to reach their final forms. A *T. gondii* ortholog of the malarial chloroquine resistance transporter protein (TgCRT) was found to be localized to the VAC membrane. Although the mutated version of the malarial chloroquine resistance transporter (PfCRT) has been shown to confer resistance to chloroquine treatment, its physiologic function remains poorly understood. Comparison between the related PfCRT and TgCRT proteins facilitates definition of the physiologic role of CRT proteins. In this study, we report that TgCRT plays a key role in regulating the integrity and proteolytic activity of the VAC and adjacent organelles, the secretion of invasion effectors, and parasite invasion and virulence. To relieve osmotic stress caused by VAC swelling when TgCRT is deleted, parasites repress proteolytic activities within this organelle to decrease solute accumulation, which then has secondary effects on parasite invasion. Our findings highlight a common function for PfCRT and TgCRT proteins in regulating apicomplexan parasite vacuolar size and function.

## Introduction

*Toxoplasma gondii* uses polypeptide invasion factors to efficiently invade host cells. These proteins are stored in two unique organelles in *Toxoplasma* parasites, the microneme and rhoptry. The micronemal proteins undergo a series of proteolytic cleavage steps within the parasite’s endosomal system, followed by further intramembrane cleavage and trimming on the parasite’s cell membrane before secretion [1,2]. Proper maturation and secretion of micronemal proteins are crucial for efficient invasion of parasites [3–5].

Micronemal protein maturation is regulated by several proteases. During intracellular trafficking, the micronemal proteins are first cleaved by aspartyl protease 3 (TgASP3) in a post-Golgi compartment [5]. A cathepsin L-like protease (TgCPL) was also shown to process some micronemal proteins in the endosome-like compartment (ELC) of the parasite [4]. The mature proteins then pass through the microneme and undergo further intramembrane cleavage and trimming on the parasite’s surface. A plasma membrane-bound protease, rhomboid 4 (TgROM4), is required to process at least some micronemal proteins and to release them from the cell surface. TgROM4 substrates include micronemal protein 2 (TgMIC2) and apical membrane antigen (TgAMA1) [5–8]. Subsequently, a subtilisin ortholog, TgSUB1, was shown to proteolyze some micronemal proteins into their final forms, including TgMIC2 and TgMIC2-associated protein (TgM2AP) [3]. Overall, precise control of proteolytic activities within the parasite’s endosomal system and on the plasma membrane is critical for processing parasite invasion effectors.

Among these proteases, TgCPL and TgSUB1 are both localized to the parasite’s endolysosomal system. TgCPL is located in an acidic digestive vacuole, termed the Vacuolar Compartment/Plant-like Vacuole (VAC) in *Toxoplasma* parasites [4,9]. Our previous studies showed that the genetic ablation of *TgCPL* causes defects in parasite invasion and acute virulence [4,10]. TgCPL becomes activated in the VAC and a portion of TgCPL is delivered to the juxtaposed ELC for maturation [4]. TgSUBI is a micronemal protease and contains a GPI anchor necessary for membrane association [3]. TgSUB1 was shown to be activated in a post-ER compartment and to enter the parasite’s endolysosomal system before trafficking to the microneme [11]. The deletion of *TgSUB1* led to inefficient trimming of micronemal proteins on the parasite surface, thereby leading to defects in invasion and virulence [3]. Hence, the maintenance of the integrity of the parasites’ endolysosomal system is critical to regulating the distribution and activity of endolysosomal proteases.

In addition to VAC dysfunction resulting in reduced invasion, replication, and virulence [4,10], parasites with impaired VAC function are unable to turn over autophagosomes during chronic infection and thereby cannot survive in host brain tissue [12]. Despite its importance, the VAC has not been well-characterized. Only 4 proteins have been localized to the VAC, including TgCPL, TgCPB (a cathepsin B-like protein), TgCRT (a *Toxoplasma* ortholog of chloroquine resistance transporter), and TgVP1 (a pyrophosphatase) [4,13–15]. The VAC forms an intact organelle during initial infection and subsequently fragments during intracellular replication [4]. It is unknown how parasites regulate these and other morphological changes that occur within the endolysosomal system. In a previous study, *TgCRT* expression was knocked down in type I *Toxoplasma* parasites using a tetracycline-inducible system [14], and VAC swelling was observed, suggesting that TgCRT is involved in VAC volume regulation. Fitness defects were also seen in these parasites suggesting that proper VAC morphology is essential for invasion and / or growth.

Interestingly, the swollen VAC phenotype for *TgCRT* knockdowns mirrors the enlarged digestive vacuole (DV) phenotype for chloroquine-resistant (CQR) *Plasmodium falciparum* expressing CQR-associated mutant PfCRT [16]. More recently, a L272F PfCRT mutation, along with CQR-conferring mutations, was found to increase DV volume by an additional 1 - 2 μm^3^ [17]. *In vitro* assays using purified recombinant PfCRT, reconstituted in proteoliposomes, suggest that PfCRT transports aminoquinoline drugs, basic amino acids, and perhaps oligopeptides, likely in an electrochemically coupled fashion [18,19]. With respect to drug transport, PfCRT expressed within CQR *P. falciparum* appears to exhibit higher chloroquine (CQ) transport efficiency relative to PfCRTs found in chloroquine-sensitive (CQS) strains [18–20]. These findings suggest that the PfCRT regulates the transport of key osmolytes from the *P. falciparum* DV. Unfortunately, the inability to successfully ablate the *PfCRT* gene [21] limits additional analysis of function *in vivo*.

Here, we successfully delete the *TgCRT* gene in a Type I *Toxoplasma* parasite strain by double crossover homologous recombination. The resulting mutant, Δ*crt*, displayed a severely swollen VAC and arrested VAC-ELC co-localization. Surprisingly, this aberrant organellar organization reduces transcription and translation of several proteases residing in the parasite’s endolysosomal system, altering microneme secretion and resulting in defective parasite invasion and acute virulence. We also engineer successful overexpression of wild type TgCRT constructs in yeast and show that the protein mediates CQ transport. Collectively, these findings determine a novel role for TgCRT in regulating VAC volume and maintaining endolysosomal integrity, suggest functional similarities for TgCRT and PfCRT proteins, and provide a new model system for analyzing the function of apicomplexan CRT proteins.

## Results

### 1. Deletion of *TgCRT* in *Toxoplasma gondii*

Previous studies have utilized an anhydrotetracycline-regulated system to reduce levels of expression of *TgCRT* in a Type I *Toxoplasma* RH strain [14]. However, incomplete depletion of *TgCRT* limits further characterization of its function. Here, we adopted a genetically tractable RH-derived strain, termed RHΔku80 (hereafter referred to as WT), to produce a complete *TgCRT* knockout. The RHAku80 strain represses the non-homologous end-joining DNA repair pathway to facilitate homology dependent DNA recombination [22]. Due to the increased homologous recombination efficiency, this strain has been widely used as a wild type *Toxoplasma* strain. We PCR amplified ~1.5-kb regions of the 5’-and 3’-untranscribed regions (UTRs) of *TgCRT*, and flanked both at both 5’-and 3’-ends with a bleomycin (*BLE*) resistance cassette to assemble a *TgCRT* knockout plasmid. The resulting construct was transfected into WT parasites to replace the entire *TgCRT* gene with a *BLE* resistance cassette by double crossover homologous recombination (**Fig 1A**). PCR analysis was performed to test for proper integration and detected the presence of the *BLE* cassette and the loss of *TgCRT* as shown in the scheme for the generation of the *TgCRT* knockout in Fig 1A. Amplification of ~1.6 kb fragments at both the 5’-and 3’-end integration regions (5’-and 3’-ARMs) were observed in the *TgCRT* knockout (Acrf), and a ~1.5 kb *TgCRT* coding region was also missing within Δ*crt* (**Fig 1B**).

**Figure 1.**
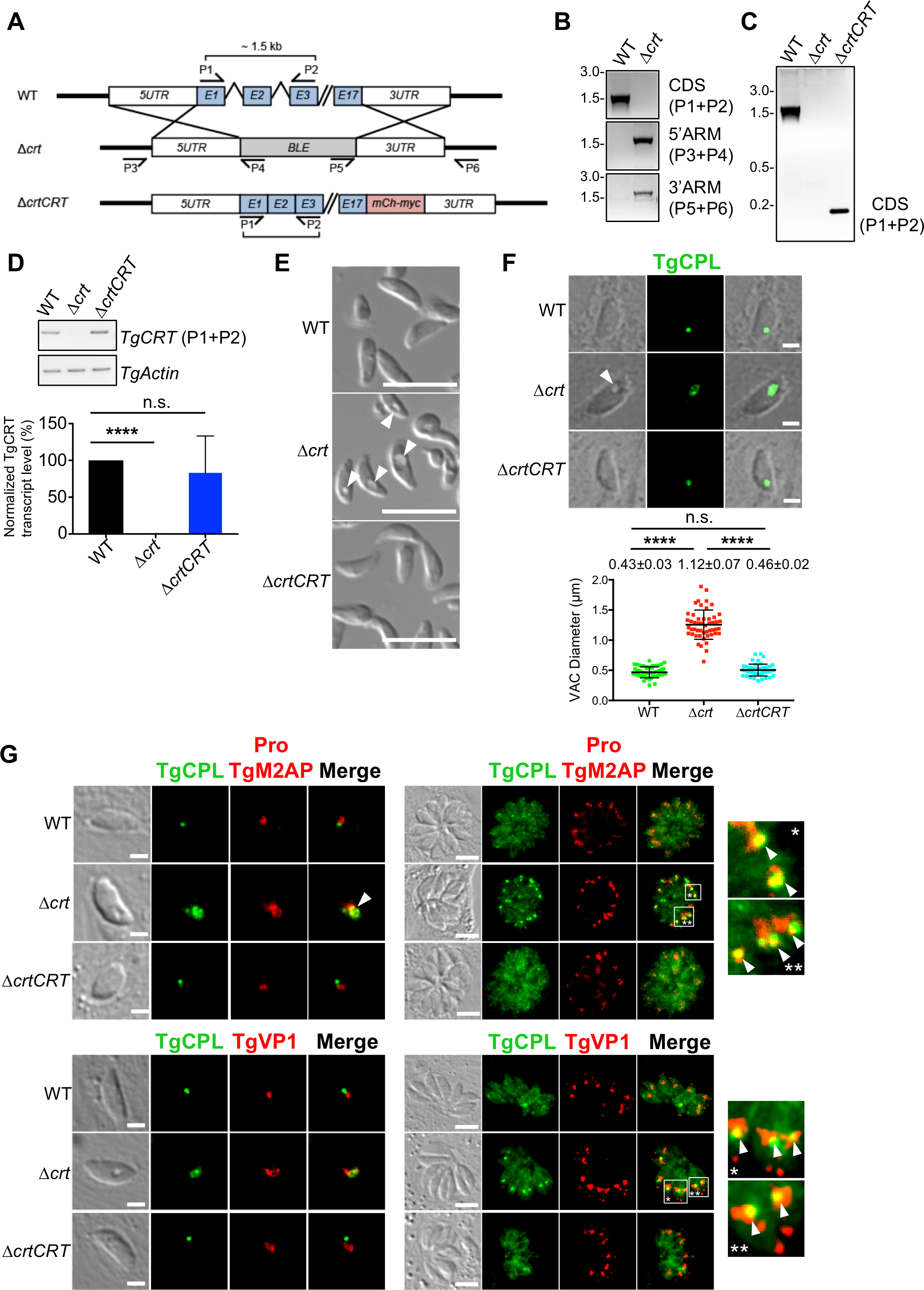
The TgCRT-deficient parasites displayed a swollen digestive vacuole and a disrupted endolysosomal system. **A)** Schematic illustration of the strategies for the *Tgcrt* deletion and complementation in *Toxoplasma* parasites. The plasmid carrying a bleomycin resistance cassette (*BLE*) flanked by the *TgCRT* targeting sequences was transfected into WT parasites for double crossover replacement of *TgCRT* to produce the Δ*crt* strain. The TgCRT complementation plasmid, containing the coding sequence of TgCRT fused with mCherry and 3xmyc epitope tags at its C-terminus, was introduced into the Δ*crt* strain to produce the Δ*crtCRT* complementation strain. **B)** The primers indicated in panel A were used to verify the correct replacement of *TgCRT* with *BLE* by PCR. **C)** The complemented *TgCRT* gene was transfected into Δ*crt* parasites and was verified by PCR. Since we complemented Δ*crt* with the coding sequence of *TgCRT*, the PCR product was a 0.2 kb fragment in the Δ*crtCRT* strain, whereas showing a 1.5 kb product in the WT strain whose *TgCRT* gene contains the introns. **D)** Transcript levels of *TgCRT* in the WT, Δ*crt*, and Δ*crtCRT* strains were evaluated by quantitative PCR. Primers were designed to anneal to the exons of *TgCRT* and are indicated in panel A. The *Tgactin* gene was included as a control for normalization. The quantification of transcripts was performed in at least three biological replicates and analyzed using unpaired Student’s *t*-test. **E)** Extracellular Δ*crt* parasites showed an enlarged concave subcellular structure, indicated by the arrow, under the differential interference contrast (DIC) microscopy. Scale bar = 5 μm. **F)** The swollen subcellular structure indicated by the arrowhead was also observed in pulse invaded Δ*crt* parasites and co-localized with a major luminal peptidase of the VAC, cathepsin L-like protease (TgCPL). The TgCPL staining was used to assess the morphology of the VAC. The parasites were allowed to invade host cells for 30 min before TgCPL antibody staining. The distance of the widest diagonal of the TgCPL staining was determined to be the diameter of the VAC and was measured using the Leica^®^ LAS X software. The measurements were conducted in three biological replicates. Measurements from 50 individual parasites from one representative assay were shown. The mean VAC size ± SD was calculated for three independent measurements and is listed on the figure. Statistical significance was determined using unpaired Student’s t-test. Scale bar = 2 μm. **G)** Parasites were costained with anti-TgCPL (the marker of the VAC) and anti-proM2AP or anti-TgVP1 (both are the markers of the endosome-like compartment, ELC). In pulse invaded parasites, the VAC and ELC staining were juxtaposed in the WT and Δ*crtCRT* strains, but aberrantly co-localized in the Δ*crt* strain. During replication, the VAC in WT and Δ*crtCRT* parasites became fragmented. However, the abnormal co-localization of the VAC and ELC significantly decreased the extent of VAC fragmentation in the Δ*crt* mutant. A major TgCPL punctum existed in the replicated Δ*crt* parasites, not in WT and Δ*crtCRT* strains, and co-localized with proTgM2AP and TgVP1 (indicated by arrows in the insets). The scale bars in the images of pulse invaded and replicated parasites are 2 μm and 5 μm, respectively. ****, *p*<0.0001; n.s., not significant.

To complement loss of *TgCRT*, we modified the pTub-TgCRT-mCherry-3xmyc plasmid (a kind gift from Dr. Giel van Dooren), which over-expresses a mCherry-3xmyc epitope-tagged *TgCRT* under control of the *Toxoplasma* tubulin promoter. A 1 kb DNA region containing the cognate *TgCRT* promoter was PCR-amplified and used to replace the tubulin promoter to provide similar transcription of complemented versus endogenous *TgCRT*. The same primer set used to detect loss of *TgCRT* in the Δ*crt* strain in **Fig 1B** was used to confirm integration of exogenously introduced *TgCRT*. Since the complemented *TgCRT* gene lacks introns, a ~0.2 kb PCR product was observed in the Δ*crtCRT* complementation strain, whereas a ~1.5 kb fragment was found for the WT strain (**Fig 1C**). To confirm that transcription of *TgCRT* in the Δ*crtCRT* strain was comparable to endogenous levels, SYBR^®^ Green-based quantitative PCR (qPCR) was used to quantify messenger *TgCRT* RNA in WT, Δ*crt*, and Δ*crtCRT* strains. No *TgCRT* transcripts were observed in Δ*crt*, further validating successful gene disruption. *TgCRT* transcript levels were similar between the WT and Δ*crtCRT* strains (**Fig 1D**). These data showed successful ablation of the *TgCRT* gene in *Toxoplasma* parasites, and properly restored expression with exogenously introduced *TgCRT* for the Δ*crtCRT* strain.

### 2. TgCRT-deficient parasites lose endolysosomal system integrity due to altered VAC morphology

Upon obtaining RHΔ*ku80*Δ*crt*, we observed that purified extracellular Δ*crt* parasites exhibited large concave subcellular structures under differential interference contrast (DIC) microscopy, whereas WT and Δ*crtCRT* strains did not display this phenotype (**Fig 1E**). This subcellular structure was also observed in pulse invaded Δ*crt* parasites (**Fig 1F**). To identify the swollen structures, we stained the WT, Δ*crt*, and Δ*crtCRT* parasites with anti-TgCPL antibodies. TgCPL is a major luminal endoprotease in the VAC of *Toxoplasma* [4,23]. Immunofluorescence microscopy showed that TgCPL staining co-localized with concave subcellular structures in Δ*crt* (**Fig 1F**). The TgCPL staining in Δ*crt* was larger than in WT and Δ*crtCRT* parasites, indicating that the VAC becomes swollen when TgCRT is absent. We quantified the VAC sizes based on TgCPL staining as described previously [12,14]. VAC diameter for the Δ*crt* parasites (1.12 ± 0.07 μm, mean ± standard deviation) is approximately 2.6-fold larger than for WT parasites (0.43 ± 0.03 μm), while for Δ*crtCRT* (0.46 ± 0.02 μm) VAC was similar to that measured for WT parasites (**Fig 1F**). If we assume the VAC is approximately spherical, then the Δ*crt* parasite VAC is approximately 15-fold larger than the WT VAC (**Fig 1F**). In contrast to pulse invaded parasites, the swollen concave structure was not observed in replicated Δ*crt* parasites (**Fig 1G**). However, TgCPL staining showed differences between WT and Δ*crt* parasites (**Fig 1G**). For WT, VAC fragments that appear during replication appear as fragmented puncta upon TgCPL staining [4]. However, TgCPL staining revealed a single punctate structure in replicating Δ*crt* parasites (**Fig 1G**). Overall, we find that loss of TgCRT severely alters the morphology of the VAC in growing as well as replicating *Toxoplasma*.

The VAC is a lysosome-like organelle, participating in the parasite’s endolysosomal system. It provides an environment for maturation of TgCPL and delivers activated TgCPL to its adjacent endosome-like compartment (ELC) to assist in processing micronemal proteins required for parasite invasion [4]. It also serves as a digestive vacuole to digest endocytosed proteins [4,10,24]. We hypothesized that the dramatic swelling of the VAC might affect the integrity of the parasite’s endolysosomal system. We stained WT, Δ*crt*, and Δ*crtCRT* parasites with antibodies recognizing markers of the VAC (anti-TgCPL) and of the ELC (anti-proTgM2AP or TgVP1) [4]. In pulse invaded parasites, the VAC and ELC displayed distinct subcellular staining in WT and Δ*crtCRT* strains, whereas in Δ*crt* parasites both markers partially co-localized (**Fig 1G**). Similarly, the single TgCPL punctum in replicating Δ*crt* parasites also showed partial co-localization with both proTgM2AP and TgVP1 (**Fig 1G**). These findings suggest that TgCRT mediated VAC morphology affects the integrity of the parasite’s endolysosomal system.

### 3. RHΔ*ku*80Δ*crt* shows reduced invasion and acute virulence

*Toxoplasma* utilizes exocytosis and endocytosis via the endolysosomes to release micronemal invasion effectors, and to ingest host proteins required for intracellular growth, respectively [10,25–27]. We therefore characterized invasion, replication, and egress for the RHΔ*ku80*Δ*crt* strain. First, we measured the invasion efficiency of parasites at 30-120 min post-infection. At 30 min post-infection, the Δ*crt* mutant showed ~50% reduction in invasion compared to WT and Δ*crtCRT* (**Fig 2A**). The differences in invasion efficiency between WT and Δ*crt* were reduced to ~20% at 60 min post-infection, and were not seen at 120 min post-infection (**Fig 2A**), suggesting that Δ*crt* parasites have slower invasion kinetics relative to the WT strain. Second, we used immunofluorescence microscopy to quantify parasite replication. Infected cells were stained with DAPI and anti-TgGRA7 antibody to define individual parasite nuclei and parasitophorous vacuolar (PV) membranes, respectively. The average number of parasites per PV was calculated for each strain in order to compare replication rates. There were no statistical differences in parasite replication between WT and Δ*crt* parasites at 28 and 40 hrs post-infection (**Fig 2B**). We also introduced NanoLuc^®^ luciferase into WT, Δ*crt*, and Δ*crtCRT* parasites, and measured the fold-change in luciferase activity for 72 hrs post-infection in order to calculate relative growth rates. Similarly, we did not observe growth differences between WT and Δ*crt* at 24, 48, and 72 hrs post-infection (**Fig 2B**). Third, the egress efficiency of each strain was determined by a lactate dehydrogenase release-based assay. The parasites were incubated with 0.5 mM Zaprinast for 5 min to induce egress. The egressed parasites disrupt host cell membranes to release lactate dehydrogenase that is subsequently quantified to extrapolate to the number of egressed PVs. We did not observe egress defects in the TgCRT-deficient parasites (**Fig 2C**). Last, we determined the acute virulence of Δ*crt* parasites in a murine model. Outbred CD-1 mice were infected with a subcutaneous or intravenous inoculum of 100 WT, Δ*crt*, or Δ*crtCRT* parasites. Thirty percent of mice infected with the Δ*crt* mutant survived when mice were infected by subcutaneous injection, while the WT and Δ*crtCRT* infections led to mortality at 10-12 days post-infection (**Fig 2D**). Mice receiving WT parasites by intravenous inoculation showed mortality starting at 13 days post-infection and all were expired at 20 days. The Δ*crt* and Δ*crtCRT* parasites caused death in 40% and 80% of infected mice respectively (**Fig 2D**). Statistical analysis showed that mice infected with WT and Δ*crt* parasites have significantly different survival. Seroconversion of the surviving mice was confirmed by ELISA (not shown). We also challenged surviving Δ*crt* mice with 1000 WT parasites by subcutaneous injection, and did not observe lethality after 30 days post-challenge. These findings indicate that the pre-inoculation of Δ*crt* parasites conferred immunological protection to subsequent acute toxoplasmosis. Collectively, our findings revealed that *Toxoplasma* parasites require the TgCRT protein for optimal invasion and acute virulence but not for replication and egress.

**Figure 2.**
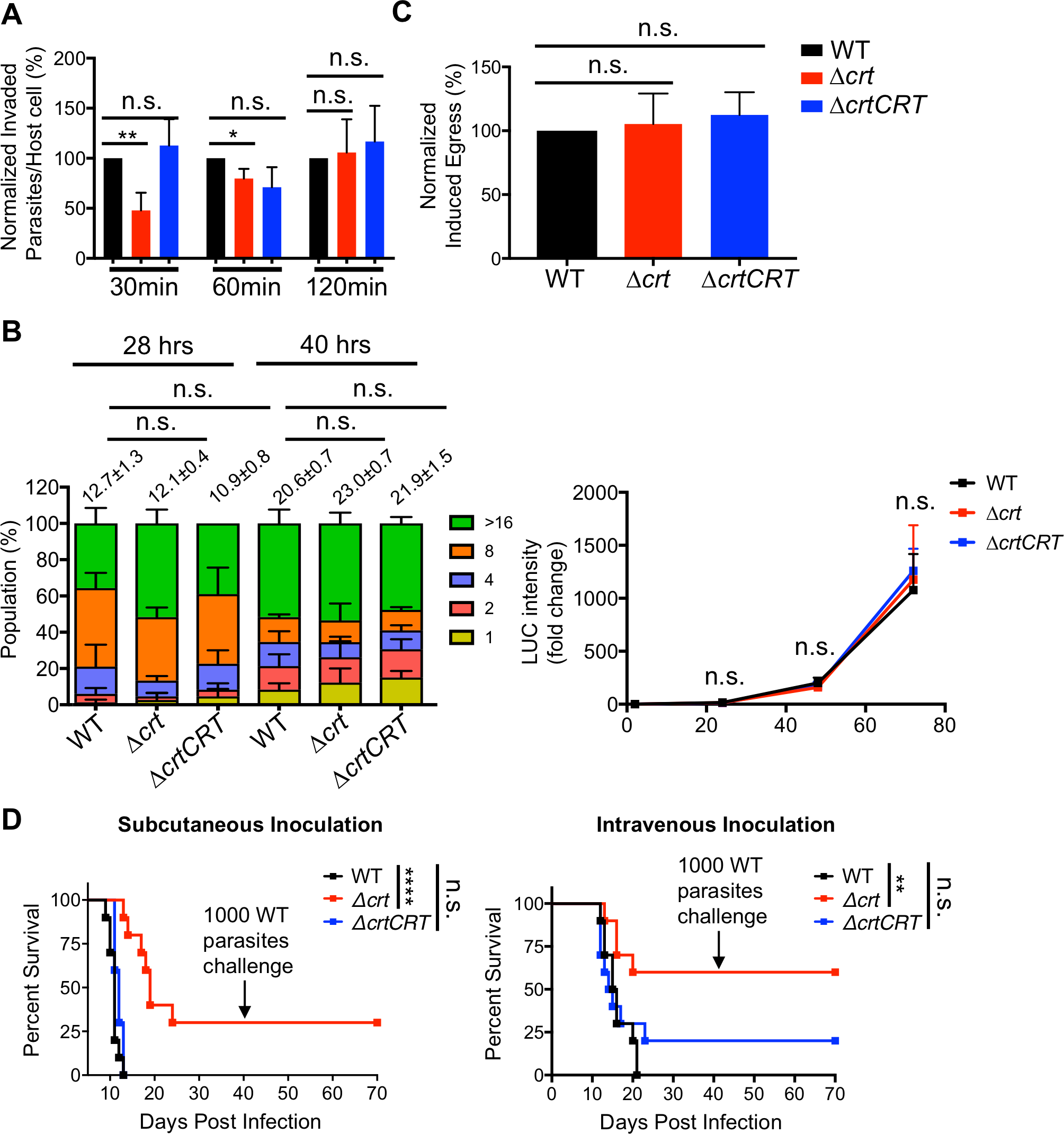
Parasite invasion and acute virulence were reduced in the Δ*crt* parasites. **A)** WT, Δ*crt*, and Δ*crtCRT* parasites were allowed to invade host cells for 30, 60, and 120 min prior to fixation and antibody staining. Parasites and host cells within six fields of view were counted for each strain. At least three independent invasion assays were conducted for statistical analysis. The Δ*crt* parasites showed a ~50% reduction in invasion at 30 min post-infection. The extent of the invasion defect in the Δ*crt* mutant was gradually minimized overtime. At 60 min post-infection, there was approximately a 25% reduction in invasion in Δ*crt* mutant, while there were no significant differences among the three strains at 120 min post-infection. The assay was performed at least in triplicate. Statistical significance was determined using unpaired Student’s t-test. **B)** We infected confluent HFFs with WT, Δ*crt*, and Δ*crtCRT* parasites for 28 and 40 hrs before fixation and staining. Infected cells were stained with DAPI and anti-TgGRA7 antibodies in order to recognize individual parasites and parasitophorous vacuoles, respectively. One hundred parasitophorous vacuoles (PVs) were enumerated for the number of parasites they contained and the distribution of different sized PVs was plotted. The average number of parasites per PV was also calculated and listed above the plots. In addition, the growth rates of these strains were determined at 24, 48, and 72 hours by using a luminescence-based assay. Parasites were inoculated into a 96-well plate pre-seeded with HFFs prior to lysis and quantification of luminescence activities at pre-determined time intervals. At least three biological replicates were performed for the replication assay. Unpaired Student’s t-test was performed to calculate the statistical significance of parasite growth between strains. **C)** To measure the egress of the parasites, 5 × 10^4^ parasites were used to infect confluent HFFs in a 96-well plate for 24 hrs. Replicated parasites were treated with 500 μM Zaprinast for 5 min at 37°C and 5% CO_2_ to induce egress. Disruption of host cell membranes due to parasite egress released lactate dehydrogenase into the medium, which was quantified and plotted. Unpaired Student’s t-test was used to calculate the statistical significance. There were no differences observed in parasite egress between WT and Δ*crt* parasites. **D)** The acute virulence of TgCRT-deficient parasites was evaluated in a murine model via subcutaneous and intravenous infections. One hundred parasites from each strain were used to infect outbred CD-1 mice (n=10 mice for each strain). The mortality of the mice was monitored for 30 days. Seroconversion of the surviving mice was evaluated by ELISA to confirm successful infection. Additionally, the surviving mice were allowed to rest for 10 days before subsequent challenge with 1,000 WT parasites by subcutaneous inoculation. The Δ*crt* mutant exhibited reduced acute virulence compared to the WT and Δ*crtCRT* strains and conferred immunological protection in the surviving mice. Data were recorded and are presented using the Kaplan-Meier plot. Statistical analysis was performed using the Log-rank (Mantel-Cox) test. For all statistical significance calculation, *, *p*<0.05; **, *p*<0.01; ****, *p*<0.0001; n.s, not significant.

### 4. RHΔ*ku80*Δ*crt* shows impaired microneme secretion

During infection, *Toxoplasma* parasites sequentially secrete proteins to facilitate host invasion. Micronemal proteins are the first to be secreted. These traffic through the parasite’s endolysosomal system and undergo intracellular maturation, intramembrane cleavage, and cell surface trimming before secretion [3–8,28]. To test which step(s) was (were) affected in the Δ*crt* parasites, we probed cell lysates and excretory secretory antigen (ESAs) fractions of each strain with anti-TgMIC2, anti-TgM2AP, and anti-TgMIC5 (antibodies to three representative micronemal proteins) by immunoblotting to measure abundances and secretion patterns. The migration patterns of these three micronemal proteins in cell lysates were very similar among the strains. The abundances of the individual micronemal proteins were normalized against the protein level of *Toxoplasma* actin protein by densitometry and plotted for quantification. All three strains showed comparable steady state abundances of these proteins (**Fig 3A**). To further evaluate abundances, we probed constitutive and induced ESAs with the same antibodies. The constitutive and induced ESAs were generated by incubating purified parasites in D10 medium (DMEM medium supplemented with 10% (v/v) cosmic calf serum) for 30 min at 37°C or D10 medium supplemented with 1% (v/v) ethanol for 2 min at 37°C, respectively. In the ESAs secreted by WT parasites, TgMIC2 exhibited two bands migrating at 100 kDa and 95 kDa, while TgM2AP showed 4 proteolytically processed polypeptides along with pro-and mature forms. However, TgMIC2 only existed as a 100 kDa band in Δ*crt* parasites. Furthermore, mature TgM2AP was not processed in the constitutive ESAs of the Δ*crt* strain and showed significantly reduced processing in the induced ESAs of Δ*crt* parasites (**Fig 3B**). The secreted TgMIC5 protein displayed similar migration patterns among these strains (**Fig 3B**). Secretion of these micronemal proteins was also quantified by normalizing the relative abundances of the proteins against the protein level of secreted TgGRA7, a dense granule protein. The secretion of TgMIC2, TgM2AP, and TgMIC5 were reduced by ~80%, 35%, and 40%, respectively, in the induced ESAs of Δ*crt* parasites, compared to the WT strain. The differences in the amount of microneme secretion were less significant in the constitutive ESAs. Only TgMIC2 secretion was decreased by ~55% in Δ*crt* parasites compared to the WT strain, while TgM2AP and TgMIC5 did not show differences (**Fig 3B**). To examine whether the abnormal secretion of micronemal proteins alters their intracellular trafficking patterns, we stained pulse invaded and replicated parasites with TgMIC2 and TgM2AP antibodies. Both microneme proteins trafficked to the apical end of the parasites and showed normal staining patterns (**Fig 3C**).

**Figure 3.**
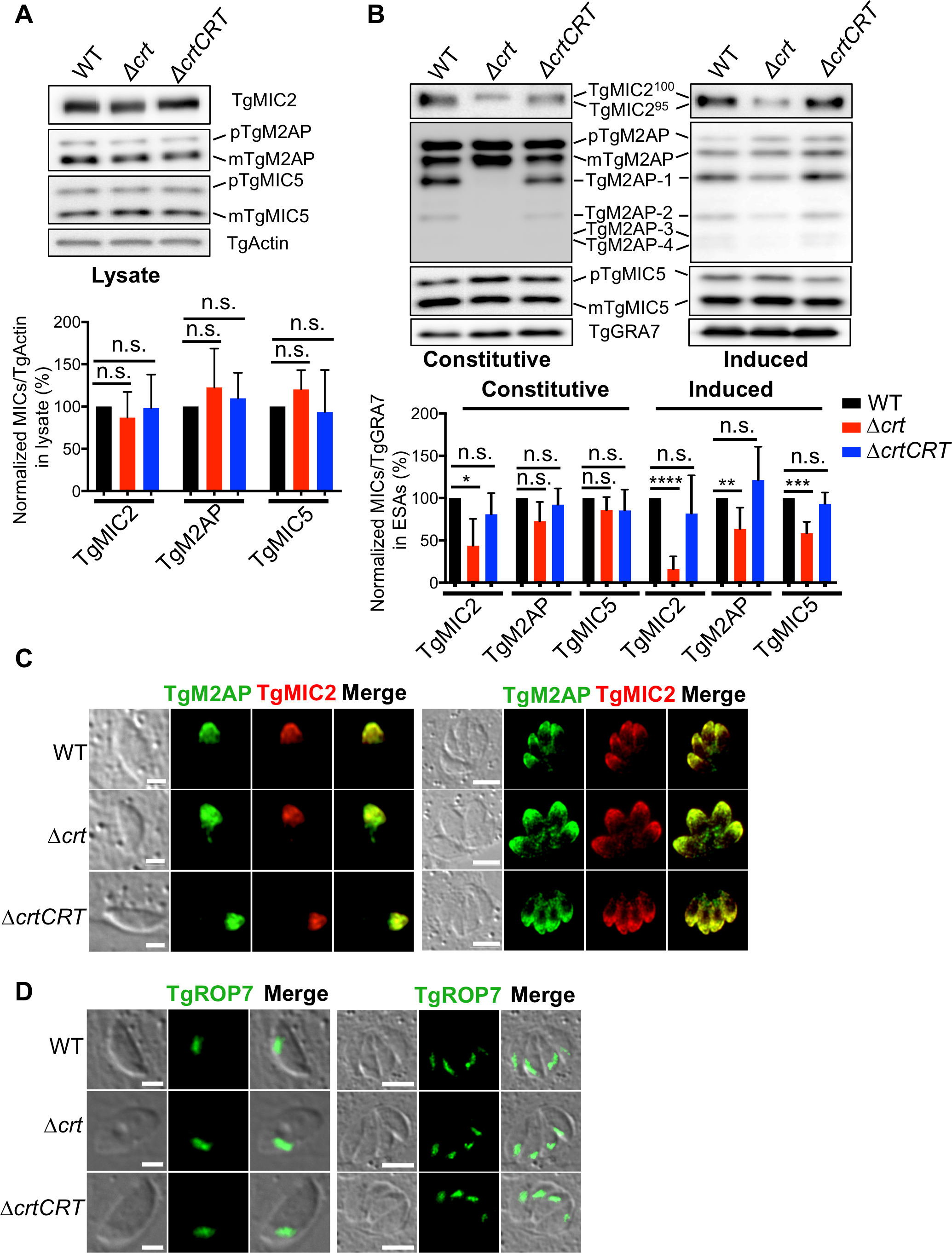
The deletion of *TgCRT* altered microneme secretion, without affecting the microneme steady abundance, intracellular trafficking, and intramembrane cleavage on the parasite surface. **A)** The steady level of micronemal proteins was not altered in the Δ*crt* parasites. Freshly lysed parasites were filter-purified, lysed, and subjected to SDS-PAGE electrophoresis and immunoblotting. The blots were probed with anti-TgMIC2, TgM2AP, and TgMIC5 antibodies, along with anti-TgActin as the loading control. Individual micronemal proteins were normalized against the corresponding TgActin to quantify their steady state expression. **B)** Δ*crt* parasites secreted less micronemal proteins than WT and Δ*crtCRT* parasites, and altered the micronemal secretion patterns. Freshly filter-purified parasites were incubated in medium at 37°C for 30 min to make constitutive ESAs, or were treated with 1% (v/v) ethanol in medium to produce induced ESAs. The ESA fractions were separated and probed with anti-TgMIC2, TgM2AP, and TgMIC5 antibodies for quantification of the secreted forms of these micronemal proteins. The ESA fractions were also probed with anti-TgGRA7 antibody, a dense granule protein, as the loading control. Statistical significance was determined using unpaired Student’s t-test. **C)** Pulse invaded and replicated parasites were stained with anti-TgMIC2 and anti-TgM2AP antibodies to examine their intracellular trafficking. No defects were detected in their intracellular trafficking. **D)** Pulse invaded and replicated parasites were also stained with anti-TgROP7 to examine the morphology of the rhoptry and intracellular trafficking of TgROP7. The rhoptry kept similar morphology and trafficking patterns among WT, Δ*crt*, and Δ*crtCRT* strains. The scale bars in the images of pulse invaded and extracellular parasites are 2 pm, and the scale bar in the images of replicated parasites is 5 μm. *, *p*<0.05; **, *p*<0.01; ***, *p*<0.001; ****, *p*<0.0001; n.s., not significant.

Prior to secretion, some membrane-anchored micronemal proteins are released via proteolytic cleavage by intramembrane rhomboid proteases such as TgROM4. The deletion of TgROM4 leads to retention of some micronemal proteins on the parasite’s plasma membrane, such as TgMIC2 and TgAMA1 (*Toxoplasma* apical membrane antigen 1) [6–8,28]. To test whether the aberrant endolysosomal system alters the retention of micronemal proteins on the surface of parasites, we stained the purified, non-permeabilized extracellular parasites with anti-TgMIC2 antibody. Immunofluorescence microscopy did not reveal excess TgMIC2 on the plasma membrane of Δ*crt* parasites (**Fig S1**), suggesting that the reduced secretion of micronemal proteins is not due to inefficient intramembrane cleavage of micronemal proteins on the parasite’s plasma membrane.

The endosome-like compartment is involved not only in the trafficking of micronemal proteins, but also rhoptry contents [29]. We stained newly invaded and replicated parasites with anti-TgROP7 antibodies to examine the trafficking of rhoptry proteins and the morphology of the rhoptry. The TgROP7 staining revealed typical rhoptry patterns located at the apical end of the parasites (**Fig 3D**), excluding the possibility of aberrant trafficking of rhoptry contents and possible defects in biogenesis. Taken together, our data suggest that invasion defects for Δ*crt* parasites are caused by incomplete trimming and consequent inefficient secretion of micronemal proteins, but not by altered intracellular maturation, trafficking, or intramembrane cleavage of micronemal proteins, nor by altered rhoptry morphology.

### 5. TgSUB1 transcript and protein levels are decreased for Δ*crt* parasites

The inefficient proteolytic processing of TgMIC2 and TgM2AP in RHΔ*ku80*Δ*crt* ESAs led us to investigate the abundance of *Toxoplasma* subtilisin 1 (TgSUB1) in the Δ*crt* parasites. A previous publication reported that parasites lacking TgSUB1 showed defective trimming patterns for secreted micronemal proteins, such as TgMIC2 and TgM2AP [3], which seemed, to us, to be similar to the secretion patterns observed for the Δ*crt* mutant. Therefore, we quantified secreted TgSUB1 in both constitutive and induced ESAs by probing them with an anti-SUB1 antibody, previously found to specifically react against TgSUB1 and PfSUB1 [30]. Immunoblotting analysis revealed that there was no detectable TgSUB1 in the ESAs of Δ*crt* parasites (**Fig 4A**). Non-permeabilized extracellular parasites were also stained with anti-SUB1 to evaluate the amount of surface-anchored TgSUB1. Similarly, there was no detectable TgSUB1 staining on the plasma membrane of Δ*crt* parasites (**Fig 4B**). These data suggest that TgSUB1 is not efficiently delivered to the surface of parasites in the Δ*crt* mutant.

**Figure 4.**
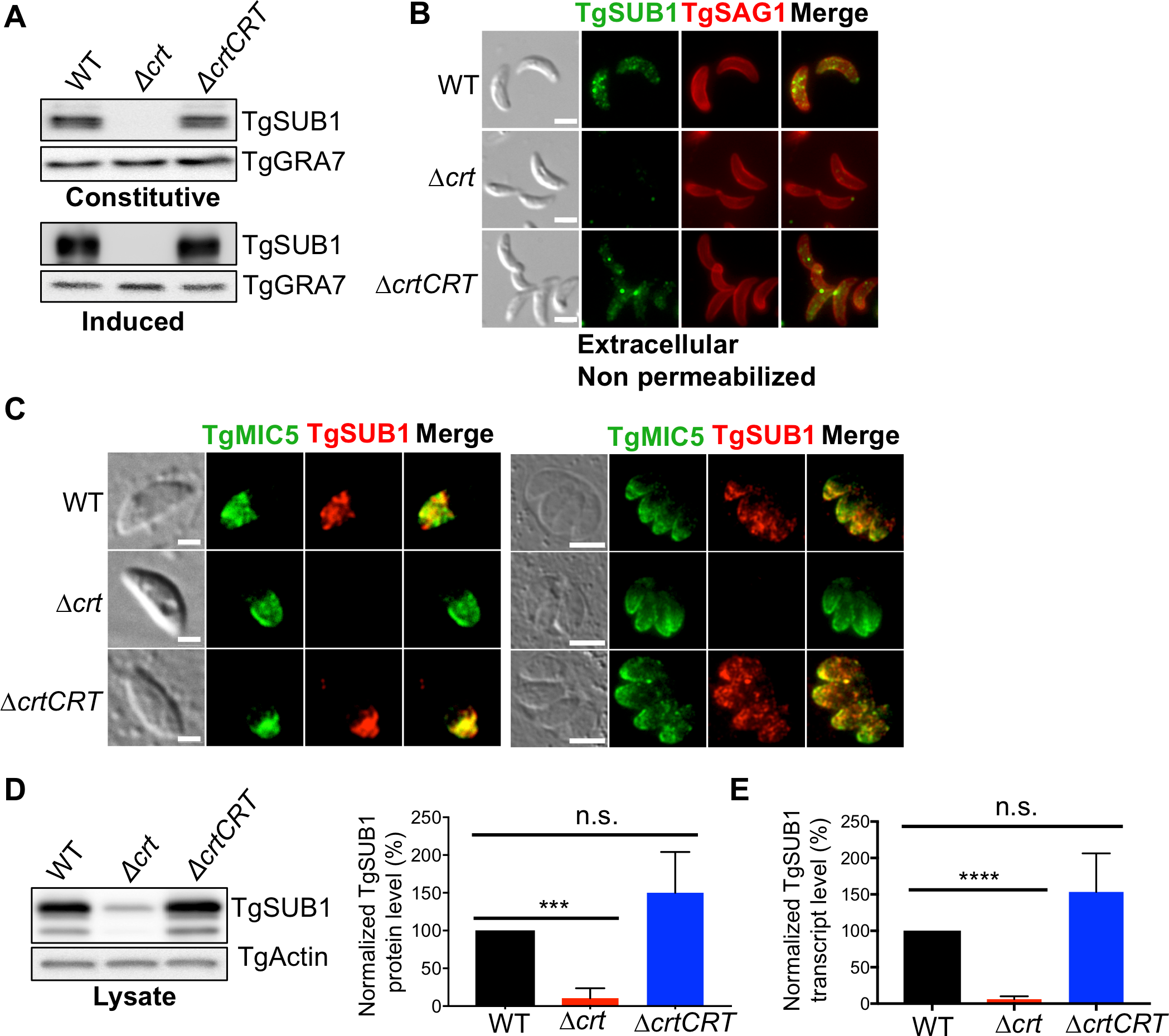
The transcript and protein levels of a subtilisin-related protease, TgSUBI, were reduced in Δ*crt* parasites. **A)** The abundance of secreted TgSUB1 in the constitutive and induced ESAs was measured by immunoblotting. There was no detectable TgSUB1 in the constitutive and induced ESAs secreted by Δ*crt* parasites via immunoblotting. B) The abundance of plasma membrane-anchored TgSUB1 was measured by probing non-permeabilized parasites with antibodies recognizing TgSUB1. The Δ*crt* mutant significantly reduced TgSUB1 on its plasma membrane. C) TgSUB1 is a micronemal protein. TgSUB1 and TgMIC5 were both found to localize in the microneme of pulse invaded and replicated WT and Δ*crtCRT* parasites. However, TgSUB1 signal was not detected in the Δ*crt* parasites. D) The cell lysates of the Δ*crt* parasites showed that the steady level of TgSUB1 was reduced by ~90% in the Δ*crt* strain. E) The transcript level of TgSUB1 was quantified by quantitative PCR. It was also decreased by approximately 90% in the Δ*crt* mutant. All assays listed in this figure were replicated at least in triplicate. Statistical significance was calculated using unpaired Student’s t-test: ***, *p*<0.001; ****, *p*<0.0001; n.s., not significant.

TgSUB1 is a micronemal protein that also traffics through the parasite’s endolysosomal system [3,11]. The aberrant endolysosomal system in Δ*crt* parasites potentially alters intracellular trafficking and/or maturation of TgSUB1 that then reduces expression. To test these possibilities, first, we stained pulse invaded and replicated parasites with anti-SUB1 to examine TgSUB1 intracellular trafficking patterns. TgMIC5 localization was used as a reference for typical expected microneme staining. Surprisingly, we observed much less TgSUB1 staining in Δ*crt* parasites compared to the WT strain (**Fig 4C**). Next, we quantified abundance of TgSUB1 in parasite cell lysates and found that TgSUB1 was decreased by approximately 90% in Δ*crt* parasites compared to WT parasites (**Fig 4D**). To further understand how TgSUB1 expression is suppressed in the Δ*crt* mutant, we performed qPCR to measure *TgSUB1* mRNA for WT, Δ*crt*, and Δ*crtCRT* parasites. *TgSUB1* transcript was reduced ~10-fold upon deletion of *TgCRT* (**Fig 4E**). Collectively, our findings suggest that arrested co-localization of the VAC and ELC dramatically decreases the abundance of TgSUB1 protein, which then alters the proteolytic processing of normally secreted micronemal protein invasion effectors, thereby reducing invasion efficiency.

### 6. VAC alterations reduce endolysosomal protease proteins and transcripts

The swollen VAC and its aberrant co-localization with the ELC in the Δ*crt* parasites could conceivably lead to altered gene transcription to assist in the adaptation of these parasites. We conducted transcriptome sequencing to detect global alterations in gene transcription for Δ*crt* parasites relative to WT. The differential gene expression analysis identified 102 genes whose transcript levels changed greater than 1.5-fold in the Δ*crt* strain. Forty-six and fifty-six genes had increased and reduced transcripts, respectively (**Fig 5A** **and Table S1**).

**Figure 5.**
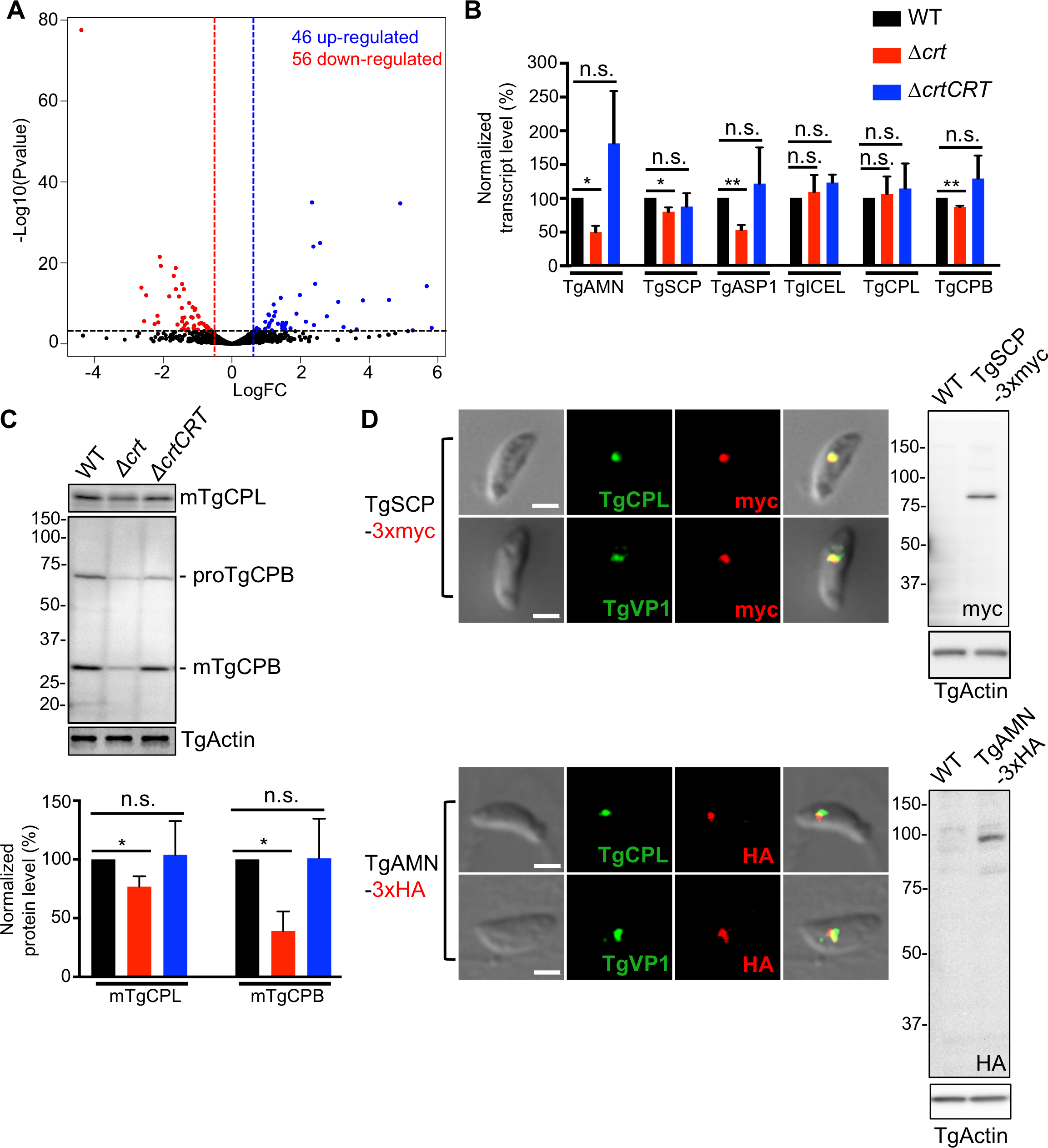
The transcript and protein abundances of several VAC-residing proteases were decreased in Δ*crt*. **A)** RNA-Seq was performed in WT and Δ*crt* parasites. Each sample was sequenced in duplicate for statistical comparison. A volcano plot was used to summarize genes that altered their transcription greater than 1.5-fold with statistical significance less than 0.05 in the Δ*crt* mutant relative to the WT strain. Forty-six and fifty-six genes labeled in the blue and red dots became up-and down-regulated in the Δ*crt* mutant, respectively. The blue and red dash lines represented the borderline of 1.5-fold change in gene transcripts, and the genes above the black dash line showed their *p* values of statistical significance below 0.05. B) qPCR was used to validate 4 down-regulated proteases that were identified by RNA-Seq analysis, along with two known VAC proteases, TgCPL and TgCPB. *TgAMN, TgSCP, TgASPI*, and *TgCPB* displayed down-regulated transcription in Δ*crt* parasites compared to WT. C) The steady protein abundances of TgCPL and TgCPB were quantified in the lysates of parasites by immunoblotting. The protein levels of TgCPL and TgCPB in Δ*crt* mutant were reduced by ~25% and 60%, respectively, compared to the WT parasites. D) TgSCP and TgAMN were endogenously tagged with 3xmyc and 3xHA, respectively, at their C-termini. The expression of the epitope-tagged proteins was confirmed by immunoblotting. The parasites were co-stained with antibodies recognizing their respective epitope tags as well as the VAC and ELC markers. Both TgSCP and TgAMN were localized in the VAC and ELC by immunofluorescence. Statistical significance was performed using unpaired Student’s *t*-test. *, *p*<0.05; **, *p*<0.01; n.s., not significant.

Four proteases were among the list of genes showing reduced transcripts in the Δ*crt* mutant, including one putative aminopeptidase N protein (TgAMN, TGGT1_221310), one putative Pro-Xaa serine carboxypeptidase (TgSCP, TGGT1_254010), aspartyl protease 1 (TgASP1, TGGT1_201840), and an ICE family protease-like protein (TgICEL, TGGT1_243298). We validated transcript levels for these proteases, as well as two known VAC luminal proteases (TgCPL and TgCPB), in WT, Δ*crt*, and Δ*crtCRT* strains by qPCR. The qPCR analysis showed that the transcript levels of TgAMN, TgSCP, TgASP1, and TgCPB were decreased by 50%, 20%, 47%, and 14%, respectively, in Δ*crt* parasites (**Fig 5B**). Protein levels of TgCPL and TgCPB were quantified by immunoblotting and compared for WT, Δ*crt*, and Δ*crtCRT* parasites. Although TgCPL transcript levels did not differ, abundance of TgCPL protein was decreased ~25% in the Δ*crt* mutant (**Fig 5C**). *TgCPB* expression was reduced and both the pro-and mature forms of TgCPB protein were reduced relative to WT parasites. (**Fig 5C**).

To determine the subcellular locations of these down-regulated proteases, we tagged endogenous TgAMN and TgSCP with 3xHA and 3xmyc epitope tags at their C-termini in WT parasites, respectively (**Fig S2**). After drug-selection, we probed cell lysates from these tagged strains with anti-HA and anti-myc antibodies, respectively, to test expression. Immunoblotting revealed that the observed molecular mass of both proteins was similar to the predicted size based on primary sequences (**Fig 5D**). Next, the tagged strains were co-stained with antibodies recognizing the epitope tags along with anti-TgCPL and anti-VP1 antibodies to determine subcellular location. Immunofluorescence microscopy revealed both TgSCP and TgAMN to be in the VAC/ELC (**Fig 5D**) TgASP1 subcellular location was also determined to be within the VAC (data not shown; Dou, Z. *et al*., in preparation). Collectively, these data suggest that the swollen VAC in Δ*crt* parasites causes reduced transcription and translation of several endolysosomal proteases.

### 7. Suppression of proteolysis within the swollen Δ*crt* VAC partially restores VAC size, organellar separation, and invasion

We suspected that inhibition of proteolysis might reduce the size of the swelled VAC. We tested this hypothesis by chemically and genetically suppressing VAC proteolysis. First, we treated WT, Δ*crt*, and Δ*crtCRT* parasites with 1 μM LHVS, an irreversible inhibitor of TgCPL protease [23]. As mentioned TgCPL is a major endopeptidase involved in the maturation of micronemal proteins and digestion of host proteins [4,10]. Infected host cells were incubated with LHVS for 48 hrs to allow full inhibition of TgCPL. Treated parasites were liberated from host cells and used to infect new host cells for 30 minutes, followed by TgCPL staining to quantify the size of the VAC. As expected, LHVS-treated Δ*crt* parasites displayed smaller VACs than DMSO-treated Δ*crt* parasites (**Fig 6A**). TgCPB is another known VAC-localizing protease, displaying both endo-and exo-peptidase activities [13,23]. Due to its carboxypeptidase activity, it is expected that TgCPB generates more small solutes relative to TgCPL. We used CRISPR-Cas9 editing to generate a Δ*crt*Δ*cpb* double knockout (**Fig 6B**). The replacement of *TgCPB* with a pyrimethamine resistance cassette was confirmed by PCR and immunoblotting (**Fig 6B**). The resulting Δ*crt*Δ*cpb* mutant showed a smaller concave subcellular structure compared to the Δ*crt* mutant (**Fig S3**). The size of the VAC in WT, Δ*crt*, and Δ*crt*Δ*cpb* was quantified based on the TgCPL staining as described above and the Δ*crt*Δ*cpb* parasite VAC (0.85 ± 0.15 μm) was reduced by ~30% compared to Δ*crt* parasites (1.15 ± 0.10 μm) (**Fig 6C**). The moderate decrease in the size of the VAC in the Δ*crt*Δ*cpb* strain also reduced the number of parasites showing partial overlap between the VAC and ELC. Approximately 44% of Δ*crt*Δ*cpb* parasites showed partial overlap between TgCPL and proTgM2AP staining compared to 62% in the Δ*crt* strain (**Fig 6D**), with both significantly higher than the 19% and 25% seen for WT and Δ*crtCRT* strains, respectively. TgSUB1 showed comparable expression in both the WT and Δ*crt*Δ*cpb* strains (**Fig 6E**). Similarly, TgSUB1 was observed in both constitutive and induced ESA fractions in the Δ*crt*Δ*cpb* parasites (**Fig 6F**). TgM2AP and TgMIC2 were cleaved by TgSUB1 in Δ*crt*Δ*cpb*, and their secretion patterns were similar to those seen in the WT strain (**Fig 6F**). These partially restored phenotypes in the Δ*crt*Δ*cpb* mutant improved the invasion efficiency by ~ 60% compared to the Δ*crt* strain, although invasion was still significantly lower than that of WT parasites (**Fig 6G**). Collectively, these data show a close association between the size of the VAC, altered morphology of the parasite’s endolysosomal system, protein abundance of TgSUB1, and parasite invasion.

**Figure 6.**
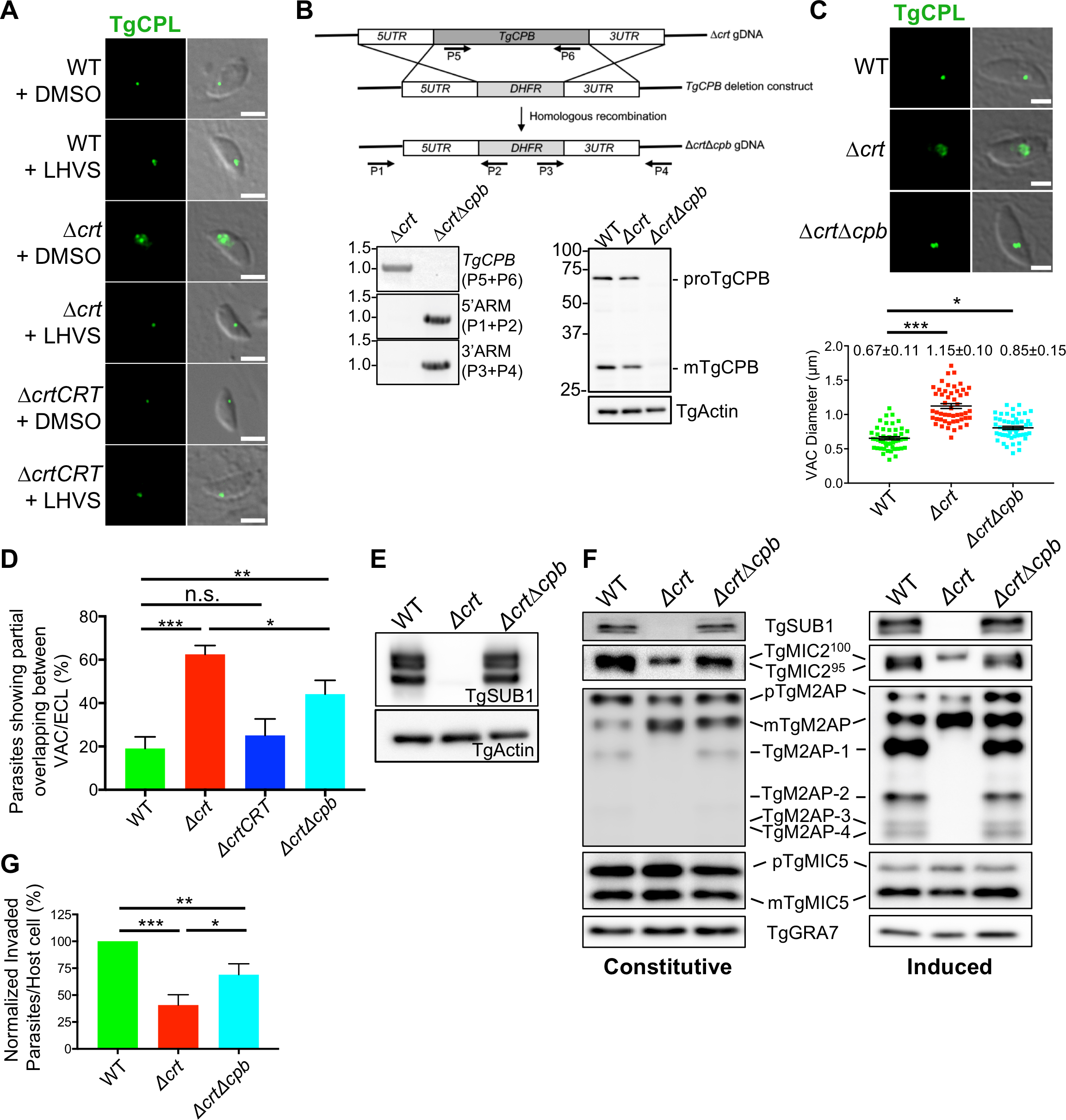
Suppression of proteolysis in the VAC reduced VAC size and partially restored integrity of parasite’s endolysosomal system, which recovered TgSUBI expression, micronemal protein trimming in ESAs, and parasite invasion. **A)** The Δ*crt* parasites were incubated with 1 pM LHVS, an irreversible inhibitor of TgCPL, for one lytic cycle, followed by a pulse invasion. Parasites were stained with anti-TgCPL antibodies to determine the size of the VAC. The swollen VAC phenotype was significantly reduced in the LHVS-treated parasites. Scale bar = 2μm. **B)** TgCPB, a VAC-residing endo-and exopeptidase was genetically deleted in the Δ*crt* strain. Schematic illustration for the creation of the Δ*crt*Δ*cpb* mutant. A PCR product carrying a pyrimethamine resistance cassette (DHFR) flanked by 50 bps of the 5’-and 3’-untranscribed regions of *TgCPB* was transfected into WT parasites for double crossover replacement of *TgCPB*. Primers indicated in panel B were used to verify the replacement of *TgCPB* with *DHFR* via PCR and agarose gel electrophoresis. The ablation of *TgCPB* in Δ*crt*Δ*cpb* parasites was also confirmed by immunoblotting. **C)** The sizes of the VAC in WT, Δ*crt*, and Δ*crt*Δ*cpb* parasites were determined by the methods mentioned above. The Δ*crt*Δ*cpb* parasites displayed a partial reduction in the size of the VAC compared to WT parasites. **D)** The pulse invaded parasites showing co-localization between the VAC (TgCPL) and ELC (proTgM2AP) in WT, Δ*crt, ΔcrtCRT*, and Δ*crt*Δ*cpb* strains were quantified. At least 100 parasites were quantified for each replicate in a total of three replicates. The Δ*crt*Δ*cpb* parasites had a significantly lower percentage of parasites having arrested co-localization between the VAC and ELC compared to the Δ*crt* mutant. **E)** The lysates of WT, Δ*crt*, and Δ*crt*Δ*cpb* parasites were probed with antibodies recognizing TgSUB1. The steady expression of TgSUB1 was restored in Δ*crt*Δ*cpb* parasites. **F)** The constitutive and induced ESAs of WT, Δ*crt*, and Δ*crt*Δ*cpb* strains were made and probed with the antibodies indicated in the figure. The secretion and trimming of micronemal proteins were also recovered in the Δ*crt*Δ*cpb* mutant. **G)** The invasion efficiency of WT, Δ*crt*, and Δ*crt*Δ*cpb* strains was determined using the procedures mentioned above. The Δ*crt*Δ*cpb* parasites showed increased invasion efficiency compared to the Δ*crt* strain, albeit still a lower efficiency than that of WT parasites. Scale bar = 2 μm. Statistical significance was calculated using unpaired Student’s t-test. *, *p*<0.05;; **, *p*<0.01; ***, *p*<0.001; n.s., not significant.

### 8. TgCRT is a functional transporter

Finally, we attempted to express TgCRT in *S. cerevisiae* yeast following previously described strategies for PfCRT [31,32]. Native *TgCRT* cDNA did not express well in *S. cerevisiae* yeast (**Fig S5**), however, following a previously published strategy for difficult to express PfCRT mutants [20] we created a fusion gene that replaced the 300 most N-terminal residues of the *TgCRT* sequence with 111 most N-terminal residues from *S. cerevisiae* plasma membrane ATPase (PMA), which harbors a yeast plasma membrane localization sequence (**Fig S4B and S4C**). Via alignment with PfCRT (**Fig S4A**), removing the 300 N-terminal TgCRT residues that are non-homologous to PfCRT preserves all putative transmembraneous domains and inter helical loop regions. The fusion protein was well expressed in *S. cerevisiae* (**Fig S5**). Following an approach previously described for PfCRT and PfCRT mutants [20,33] we assayed PMA-TgCRT expressing yeast for chloroquine (CQ) transport (**Fig. 7A**). Via alignment with PfCRT (**Fig S4A**), TgCRT T369 corresponds to the well-studied K76 residue within PfCRT; previously, mutation of PfCRT K76 to T has been shown to increase the efficiency of CQ transport by PfCRT [20,34,35]. We individually expressed both WT TgCRT and a T369K variant in the yeast to measure their transport efficiencies. Both the wild type protein and a TgCRT T369K mutant were found to transport CQ slower than PfCRT under similar conditions and to require higher external [CQ] (80 mM versus 16 mM for PfCRT) to achieve similar levels of transport (**Fig 7A**). These initial data have shown that mutation of the corresponding TgCRT threonine to lysine affects CQ transport similarly.

**Figure 7.**
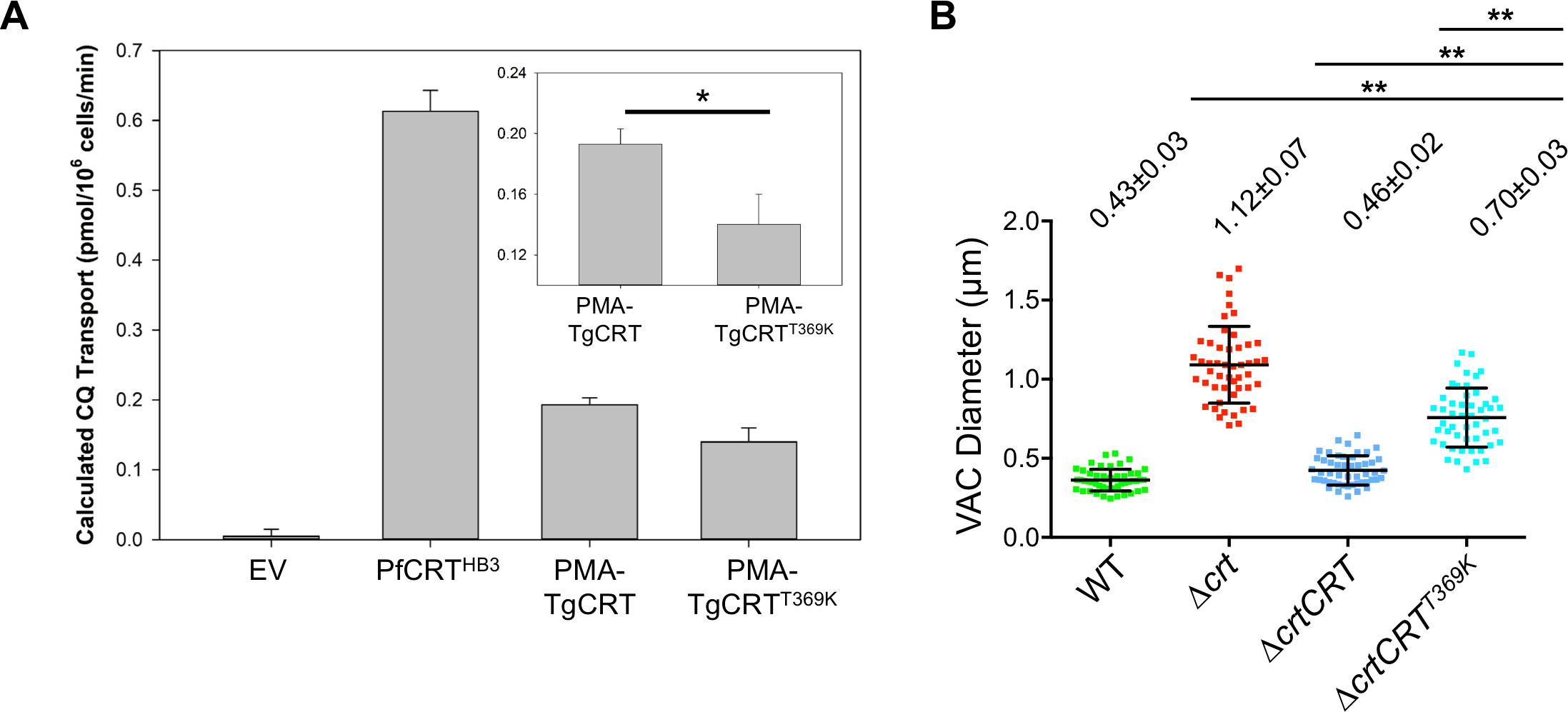
TgCRT is a functional transporter and its transport efficiency is correlated with VAC size in the parasites. **A)** CQ transport by PfCRT and TgCRT (PMA-TgCRT) expressed in *S. cerevisiae*. Transport was extrapolated from CQ-induced growth delays as described in [32,33]. Results are the average of at least four independent experiments ± SEM. EV, empty vector; PfCRT-HB3, wild type PfCRT; PMA-TgCRT, plasma membrane ATPase-TgCRT fusion (see **Fig S4**); PMA-TgCRT^T369K^, TgCRT fusion protein harboring T to K substitution at the position analogous to residue 76 in PfCRT (see text). **B)** A mutation of T369K was introduced into WT TgCRT complementation construct by site-directed mutagenesis before it was electroporated into the Δ*crt* mutant. The VAC sizes were determined based on TgCPL staining using the methods mentioned above. The Δ*crt* mutant complemented with TgCRT^T369K^ partially restored its VAC size, but it was still significantly bigger than that transfected with WT TgCRT. Statistical significance was calculated using unpaired Student’s *t*-test. *, *p*<0.05; **, *p*<0.01.

We also exchanged threonine for lysine in the WT TgCRT complementation construct and transfected Δ*crt* to examine the extent to which TgCRT^T369K^ affects VAC size in *Toxoplasma* parasites. Interestingly, in contrast to full recovery of VAC size in the WT TgCRT complementation strain, TgCRT^T369K^ only partially restored the swollen VAC (**Fig 7B**). These findings, along with the TgCRT transport data, strongly suggest that the swollen VAC is caused by luminal osmolyte excess, similar to findings for PfCRT as described in “Discussion”.

In summary, our findings strongly suggest a role for TgCRT in small solute transport that regulates VAC volume, similar to the role proposed for PfCRT [16,17]. However, at least for *T. gondii*, osmotic pressure within this organelle regulated by the TgCRT protein governs proper segregation of other organelles within the endolysosomal system that, in turn, facilitates microneme secretion and parasite invasion. The data also indicate that the invasion deficiency exhibited by the Δ*crt* mutant is likely due to multiple factors, since the recovery of TgSUB1 expression and micronemal trimming in Δ*crt*Δ*cpb* did not completely reverse invasion defects. To our best knowledge, this is the first observation of regulation of apicomplexan parasite invasion by a CRT protein.

## Discussion

*Toxoplasma* utilizes an endolysosomal system to secrete invasion effectors that disseminate infection. These invasion effectors undergo a series of intracellular proteolytic cleavage and trimming steps to reach their final forms. Therefore, maintenance of the integrity of the endolysosomal system is critical for controlling the secretion of invasion effectors in *Toxoplasma*. The Vacuolar Compartment (VAC) is an acidic lysosome-like vacuole. Previous work showed that deletion of a cathepsin L-like protease, a major VAC luminal endopeptidase, leads to invasion, replication, and virulence defects [4,10]. Compromised proteolytic activities within these parasites also result in the inefficient degradation of endocytosed host proteins [10]. Warring *et al*. previously reported that a *Toxoplasma* ortholog of chloroquine resistance transporter (TgCRT) resides in the VAC and that decreased expression of TgCRT leads to swelling of the VAC [14].

Here, we created a *TgCRT* knockout that completely removes *TgCRT* from the VAC membrane. The resulting Δ*crt* strain shows a dramatic increase in VAC size, and the organelle aberrantly co-localizes with the adjacent endosome-like compartment (**Fig 8**). Although a previous study reported that parasites deliver minor amounts of TgCPL to the ELC which then contributes to maturation of some micronemal proteins [4], our data do not reveal abnormal intracellular cleavage or trafficking of micronemal proteins in Δ*crt* parasites. Relatedly, Dogga *et al*. recently documented that aspartic acid protease 3 (TgASP3) localizes in a post-Golgi compartment and serves as a major maturase for invasion effectors [5], suggesting that the cleavage of micronemal proteins by TgCPL in the ELC plays only a minor role in their maturation.

**Figure 8.**
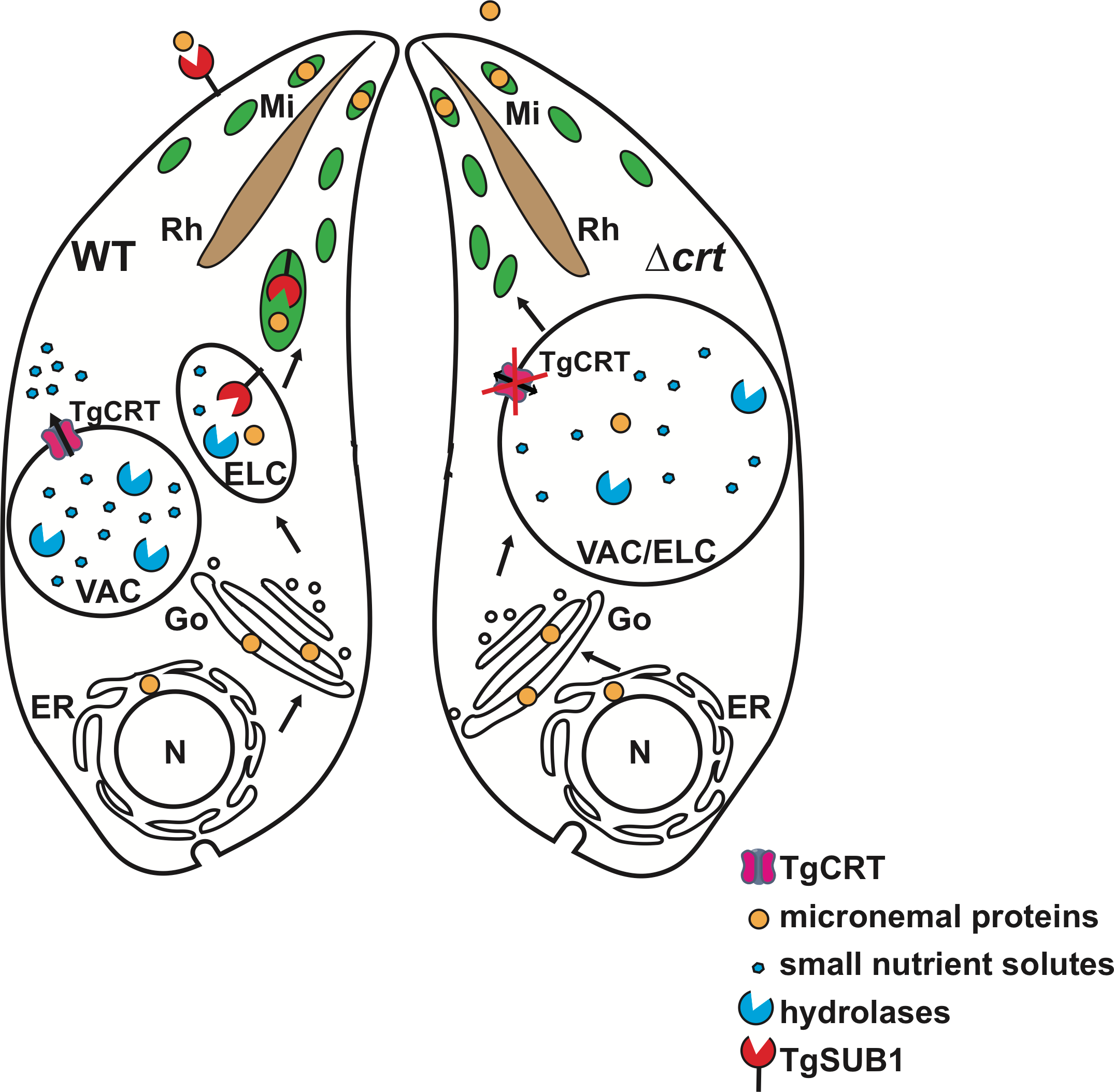
A model for the regulation of the endolysosomal system in *Toxoplasma* parasites. The *Toxoplasma* parasite contains a separate endosome-like compartment from the digestive vacuole within its endolysosomal system. When the parasite lacks TgCRT, the VAC gets swollen and cannot separate from its adjacent ELC. This aberrant co-localization leads to reduced transcript and protein abundances of several proteases residing within the endolysosomal system, including TgSUB1. These changes alter the secretion of micronemal proteins, thereby resulting in invasion defects in the TgCRT-null mutant. ELC, endosome-like compartment; ER, endoplasmic reticulum; Go, Golgi apparatus; Mi, microneme; N, nucleus; Rh, rhoptry; TgCRT, *Toxoplasma* chloroquine resistance transporter ortholog; TgSUB1, *Toxoplasma* subtilisin 1; VAC, vacuolar compartment.

We also measured the retention and secretion of micronemal proteins on the parasite’s surface and in the medium executed by TgROM4 and TgSUB1, respectively. We found that the micronemal proteins were improperly trimmed on the surface of Δ*crt* parasites. Patterns of secreted micronemal proteins observed for the Δ*crt* mutant were similar to those for Δ*sub1* parasites, which led us to examine the expression of TgSUB1 in Δ*crt* parasites and ESAs. As expected, levels of TgSUB1 were decreased on the surface of Δ*crt* parasites and in the medium during secretion. Interestingly, the steady state abundance of TgSUB1 was also significantly decreased in the Δ*crt* mutant. Surprisingly, we found that the reduction of TgSUB1 was due to a decrease in the transcription level of TgSUB1 in the Δ*crt* strain, suggesting that the parasites utilize a feedback transcriptional mechanism to regulate TgSUB1.

TgSUB1 is a micronemal GPI anchored protein. It remains unclear how TgSUB1 becomes activated within *Toxoplasma*. Previous pulse-chase experiments have revealed that TgSUB1 undergoes maturation in a post-ER compartment, and passes through the endolysosomal system before its arrival at the microneme and subsequent secretion [11]. The propeptide region of TgSUB1 carries targeting information which helps to guide the protein to the microneme [36]. The propeptide may also function by binding to active sites of mature TgSUB1 to inhibit its proteolytic activity during trafficking. Co-localization of the VAC and ELC could bring propeptide-bound TgSUB1 to a protease-abundant environment, where non-specific digestion of the propeptide could then lead to increased digestive activities in the VAC and ultimately result in an increase in osmotic pressure within the hybrid VAC/ELC organelle (**Fig 8**). During this scenario, the parasites may utilize a feedback mechanism to repress additional expression of TgSUB1 in order to avoid further VAC swelling. Moreover, we also discovered that the Δ*crt* parasites had reduced protein and/or transcript levels of several other proteases, including two known VAC proteases, TgCPL and TgCPB. Therefore, the parasites down-regulate a number of endolysosomal-VAC proteases to suppress proteolytic activities in the swollen VAC, presumably to reduce osmotic pressure and thereby control VAC size. Among these proteases, TgSUB1 has been shown to be involved in parasite invasion and virulence defects but not replication and egress [3]. Additionally, TgCPL plays a role in parasite invasion by maturing several micronemal proteins [4]. Therefore, the invasion defects exhibited in the Δ*crt* mutant could be due to several factors.

Altered endolysosomal protease transcript levels in Δ*crt* parasites suggest that parasites repress transcription factors or enhance transcription repressors to respond to increased VAC size. RNA-Seq analysis did not reveal any changes in the AP2-family of transcription factors (data not shown). In mammalian cells, the transcription factor EB (TFEB) is a master regulator that drives gene expression for autophagy and lysosome biogenesis [37]. Search of the *Toxoplasma* genome did not reveal a TgTFEB ortholog, suggesting that these parasites may adopt an alternative strategy for regulating lysosomal gene expression. Interestingly, our differential gene expression analysis identified that the transcript levels of two zinc finger (CCCH) type motif-containing proteins, TGGT1_246200 and TGGT1_226310, were increased and decreased by 2-fold and 3-fold (**Table S1**), respectively, in the Δ*crt* mutant. The CCCH type zinc finger motif-containing protein is known to regulate the stability of mRNA [38]. For example, tristetraprolin inhibits the production of tumor necrosis factor-α in macrophages by destabilizing its mRNA via an interaction with AU-rich elements at the 3’-untranslated region [39]. Further investigation to identify transcription factor(s) and regulator(s) that govern the expression of *Toxoplasma* lysosomal genes will help elucidate how these parasites regulate the biogenesis and function of the VAC.

In this study, we have determined that TgCRT-deficient parasites have reduced expression of several endolysosomal proteases. We have also found that suppression of proteolytic activities within the swollen VAC decreases the size of the organelle. These findings, along with data verifying that TgCRT is indeed a transporter with function similar to that of PfCRT, support the idea that TgCRT functions to transport essential VAC osmolytes, similar to proposals for PfCRT [16,17,40]. Likely candidate osmolytes include ions and/or amino acids. We suggest that when TgCRT is absent on the membrane of the VAC, protein degradation products (short peptides, amino acids) likely accumulate within the VAC and increase osmotic pressure, thereby leading to the swollen phenotype. Consistent with this idea, and similar to related observations for *P. falciparum* treated with cysteine protease inhibitors [41], chemical inhibition of proteolysis via the small inhibitor LHVS dramatically reduces the size of the VAC. For *Toxoplasma*, LHVS principally targets TgCPL, but also binds to TgCPB protease [13]. Additionally, the maturation of TgCPB is dependent upon the presence of TgCPL [13]. Therefore, the treatment of LHVS blocks both of these VAC proteases. Since the cathepsin B (TgCPB) protease exhibits endo-and exo-peptidase activities, and the deletion of TgCPB had no effect on the abundance of TgCPL [13], we genetically ablated *TgCPB* to further understand its role in regulating VAC morphology. The deletion of *TgCPB* partially restored the size of the VAC, secretion patterns of micronemal proteins, and invasion defects. These results reveal for the first time that TgCPB plays an active role in contributing to proteolysis within the VAC in *Toxoplasma* parasites.

RNA-Seq analysis identified several other genes with altered transcription levels, suggesting that the parasites may utilize additional strategies to control VAC size. For example, interestingly, levels of aquaporin (TGGT1_215450) transcript were reduced in Δ*crt* parasites. Previous work showed that this aquaporin is localized to the VAC/PLV [42]. Therefore, it seems likely that Δ*crt* parasites express less aquaporin to reduce water transport into the VAC/PLV, as an additional tactic to limit VAC swelling. We also found that two putative protein phosphatase 2C (TGGT1_276920 and TGGT1_201520) transcripts are down-regulated in the Δ*crt* mutant. Both carry signal peptides, indicating endosomal trafficking. TGGT1_276920 and TGGT1_201520 are homologous to PTC3 and PTC1 in *S. cerevisiae*, respectively. Interestingly, both PTC1 and PTC3 proteins are involved in yeast osmosensory regulation. A mitogen-activated protein kinase pathway is activated when yeast cells experience hyperosmotic conditions. PTC1 and PTC3 negatively regulate this pathway [43,44]. Furthermore, PTC1 was found to control the function and morphology of the yeast vacuole, which further alters its biogenesis [45]. The dramatic change in

*Toxoplasma* VAC volume indicates induced osmotic stress in the Δ*crt* parasites. The knockout parasites appear to be utilizing a similar mechanism to suppress these protein phosphatases and enhance similar osmoregulatory signaling. We suggest similar studies for *P. falciparum* and other apicomplexan parasites that express CRT orthologs would be informative.

The phenotype of the swollen VAC in the Δ*crt* strain mirrors the enlarged digestive vacuole in chloroquine (CQ) resistant (CQR) *P. falciparum* malaria [16]. Peptidomic analysis showed that hemoglobin is not as efficiently degraded within the digestive vacuole (DV) in CQR malaria parasites [40], further suggesting that CQR mutations in PfCRT alter the physiology within the swollen digestive vacuole, thereby compromising DV proteolytic activities. *In vitro* assays utilizing recombinant PfCRT, reconstituted in proteoliposomes, have revealed that PfCRT may act as a proton gradient dependent, polyspecific nutrient exporter for small solutes including amino acids, oligopeptides, glutathione, and small drugs [18,19]. These studies also demonstrate that CQR-associated PfCRTs display altered transport efficiency relative to CQ-associated PfCRT. Our study has revealed that TgCRT mediates CQ transport similar to PfCRT. The Δ*crt* strain appears more sensitive to CQ relative to WT parasites (**Fig S6**), further suggesting that TgCRT is a functional transporter of small solutes across the membrane of the VAC. We suggest that alteration of proteolytic activities in the enlarged VAC of the Δ*crt* mutant reveals a similar scenario relative to the CQR *P. falciparum* DV. Given the similarity in components and functionality of the VAC and DV found in *Toxoplasma* and *Plasmodium*, this *Toxoplasma* TgCRT-deficient mutant should prove useful for further studying the native function of CRT orthologs from other apicomplexan parasites.

In sum, our findings reveal that the *Toxoplasma* TgCRT protein is indeed a small molecule transporter that plays an essential role in regulating the size and morphology of the VAC. Unexpectedly, this regulation maintains integrity of the parasite’s endolysosomal system, which is essential for the trafficking of invasion effectors. Co-localization of the VAC and endosome-like compartment in the *TgCRT* knockout led to a reduction in transcript and protein levels for several endolysosomal proteases. We found that blocking normal proteolysis within the swollen VAC reduced the size and partially restored the morphology of the organelle. Taken together, these findings suggest that TgCRT mediates the transport of small solutes in order to regulate VAC size and morphology. The data show that the integrity of the parasite endolysosomal system is critical for parasite virulence. We suggest that pharmaceutical modulation of the VAC could serve as a novel strategy for managing toxoplasmosis.

## Materials and Methods

### Ethical statement

This study was performed in compliance with the Public Health Service Policy on Humane Care and Use of Laboratory Animals and Association for the Assessment and Accreditation of Laboratory Animal Care guidelines. The animal protocol was approved by Clemson University’s Institutional Animal Care and Use Committee (Animal Welfare Assurance A3737-01, protocol number AUP2016-012). All efforts were made to minimize discomfort. This method of euthanasia is consistent with the recommendations of the Panel on Euthanasia of the American Veterinary Medical Association.

### Chemicals and reagents

Morpholine urea-leucyl-homophenyl-vinyl sulfone phenyl (LHVS) was kindly provided by the Bogyo lab at Stanford University. Other chemicals used in this work were analytical grade and were purchased from VWR unless otherwise indicated.

### Parasite culture

*Toxoplasma gondii* parasites were cultured in human foreskin fibroblast (HFF) cells (ATCC, SCRC-1041) in Dulbecco’s Modified Eagle Medium (DMEM) media supplemented with 10% cosmic calf serum at 37 °C with 5% CO_2_. The parasites were harvested by membrane filtration as described previously [13].

### Generation of transgenic parasites

To generate the TgCRT-deficient strain, ~1.5 kilobases (kb) of the 5’-and 3’-UTR of the *TgCRT* gene, respectively, were PCR-amplified and flanked at both ends of the bleomycin resistance cassette (BLE) to assemble a *TgCRT* deletion construct. The resulting plasmids were introduced into WT parasites by electroporation. The transfected parasites were selected with 50 μg/ml bleomycin twice, while in their extracellular stage as described previously [13]. Clones of the TgCRT-deficient parasites were isolated by limiting dilution. The correct replacement of *TgCRT* with the *BLE* cassette was confirmed by quantitative PCR (see text).

To complement Δ*crt* parasites, we modified the plasmid pTub-TgCRT-mCherry-3xmyc (a gift from the van Dooren lab), which expresses a C-terminally mCherry-3xmyc epitope-tagged TgCRT under the *Toxoplasma* tubulin promoter. The plasmid was restricted with HpaI and MfeI to remove the tubulin promoter and a segment of *TgCRT*. The remaining DNA fragment served as the backbone for subsequent Gibson assembly to incorporate a PCR amplified ~1 kb region upstream of the *Tgku80* gene, the ~1 kb fragment of the *Tgcrt* 5’-UTR region, and the removed partial *Tgcrt* coding sequence to produce the TgCRT complementation plasmid, pCRT-TgCRT-mCherry-3xmyc. The complemented TgCRT is driven by its cognate promoter to maintain physiologic similarity to native TgCRT expression in WT parasites (see text). The 1 kb region located ~6 kb upstream of the *Tgku80* gene was used to facilitate a single integration of the TgCRT complementation plasmid into this specific locus by single crossover homologous recombination. The TgCRT complementation construct was digested with SwaI restriction enzyme, gel-extracted, purified, and transfected into Δ*crt* parasites by electroporation.

To introduce NanoLuc^®^ luciferase (nLuc) into parasites, we PCR-amplified and assembled the *Tgtubulin* promoter, the coding sequence of the nLuc luciferase, and an HXG selection marker into an nLuc expression construct. The resulting plasmid was transfected into WT, Δ*crt*, and Δ*crtCRT* strains. The transfectants were selected with 25 μg/ml mycophenolic acid and 50 μg/ml xanthine. Stable populations were subjected to limiting dilution to generate individual clones of WT*::nLuc*, Δ*crt::nLuc*, and Δ*crtCRT::nLuc* and clones were confirmed via luciferase activity.

To generate the Δ*crt*Δ*cpb* mutant, the *TgCPB* gene was replaced with a pyrimethamine resistance cassette using the CRISPR-Cas9 genome editing system [46,47]. The pyrimethamine resistance cassette was PCR-amplified and flanked by 50 bp regions upstream and downstream of the start and stop codons of the *TgCPB* gene for homologous recombination. A 20 bp region located at the beginning of the coding region of the *TgCPB* gene was used to design guide RNA and replace the guide RNA targeting TgUPRT gene in the plasmid pSAG1-Cas9::UPRTsgRNA using Q5 site-directed mutagenesis (NEB). The Cas9-GFP and guide RNA constructs were co-transfected into Δ*crt* parasites with the corresponding repair PCR product. The guide RNA and Cas9 generated a gap within the *TgCPB* gene to facilitate double crossover homologous recombination. Correct gene replacement was confirmed by PCR.

To epitope-tag TgAMN, we again used CRISPR-Cas9 editing tools to modify the corresponding gene. Guide RNA recognizing the 20 bp region near the TgAMN stop codon was generated using the methods above.

The 50-bp homologous regions upstream and downstream of the stop codon of the *TgAMN* gene were cloned at the 5’-and 3’-ends of the DNA sequence containing the 3xHA epitope tag and the pyrimethamine resistance cassette, respectively, by PCR. The plasmid encoding the guide RNA targeting TgAMN and Cas9-GFP and the PCR product were co-transfected into WT parasites. The stop codon of TgAMN was replaced by the 3xHA epitope tag and pyrimethamine resistance cassette. Stable populations were generated after multiple rounds of pyrimethamine selection and TgAMN-3xHA fusion protein was confirmed by immunoblotting analysis.

TgSCP was endogenously tagged with a 3xmyc epitope tag via single crossover. An approximately 1 kb region upstream of the *TgSCP* stop codon was PCR amplified and fused in frame with a 3xmyc epitope to assemble TgSCP-3xmyc. A pyrimethamine resistance cassette was also included, the resulting plasmid was linearized and transfected into WT parasites. The correct tagging was confirmed by immunoblotting.

### Site-directed mutagenesis

Threonine 369 was mutated to lysine in the WT TgCRT complementation construct, via site directed mutagenesis according to the Q5^®^ site-directed mutagenesis procedure (NEB). Linear PCR product was phosphorylated, circularized, and transformed into *E. coli*. Correct clones were identified by direct DNA sequencing.

### Transfection of *Toxoplasma* parasites

*T. gondii* parasites were allowed to grow in HFF cells for 48 hrs at 37 °C with 5% CO_2_. Freshly egressed parasites were syringed, filter purified, and harvested in Cytomix buffer (25 mM HEPES, pH 7.6, 120 mM KCl, 10 mM K_2_HPO4/ KH_2_PO4, 5 mM MgCl_2_. 0.15 mM CaCl_2_, and 2 mM EGTA). Parasites were pelleted at 1,000 × *g* for 10 min, washed once in Cytomix buffer, and resuspended in Cytomix buffer at 2.5 × 10^7^ parasites per ml. 400 μL of parasite suspension was mixed with 20 μg DNA and 2 mM ATP/5 mM reduced glutathione to a final volume of 500 μL. The mixture was electroporated at 2 kV and 50 ohm resistance using the BTX Gemini X2 (Harvard Apparatus). Transfectants were inoculated into a T25 flask pre-seeded with confluent monolayer of HFF cells and the cells allowed to recover. Drug selection was applied 24 hrs post transfection.

### Immunofluorescence

Freshly lysed parasites were used to infect confluent HFF cells pre-seeded in an 8-well chamber slide for 1 hr (pulse invaded parasites) or 18-24 hrs (replicated parasites). The extracellular parasites were attached to chamber slides using 0.1% (w/v) poly-L-lysine. Immunofluorescence was performed as described previously [10,13]. Images were viewed and digitally captured using a Leica^®^ CCD camera equipped with a DMi8 inverted epifluorescence microscope and processed with Leica^®^ LAS X software.

### Excretory secretory antigens (ESAs) preparation

Freshly egressed parasites were syringed, filter purified, and resuspended at 5 × 10^8^ parasites/ml in D1 medium (DMEM medium supplemented with 1% FBS). 100 μL of parasite suspension was transferred to a microfuge tube and incubated at 37 °C for 30 min to prepare constitutive ESAs. To isolate induced ESAs, the parasite suspension was incubated in D1 medium supplemented with 1% ethanol for 2 min at 37°C. ESAs were separated from intact parasites by centrifugation at 1,000 x *g* for 10 min. ESA fractions were transferred to a new microfuge tube, mixed with SDS-PAGE sample loading buffer, and boiled for 5 min for immunoblotting analysis.

### SDS-PAGE and Immunoblotting

Parasite lysates and ESA fractions were prepared in 1x SDS-PAGE sample buffer and boiled for 5 min before resolving on standard SDS-PAGE gels. For immunoblotting, gels were transferred to PVDF membranes by semi-dry protein transfer methods. Blots were blocked with 5% non-fat milk and incubated with primary antibody diluted in 1 % non-fat milk. Goat anti-mouse or anti-rabbit IgG antibodies conjugated with horseradish peroxidase were used as secondary antibody. Immunoblots were developed with SuperSignal™ WestPico chemiluminescent substrate (Thermo). The chemiluminescence signals were captured using the Azure^®^ Imaging System. Bands were quantified by densitometry using LI-COR^®^ Image Studio software.

### Parasite invasion assay

The red-green invasion assay was used to measure the efficiency of parasite invasion. Freshly purified parasites were syringed, filter purified, and resuspended at 5 × 10^7^ parasites/ml in invasion medium (DMEM supplemented with 3% FBS). 200 μL of parasite resuspension was inoculated into each well of an 8-well chamber slide pre-seeded with HFF cells, and parasites were allowed to invade host cells for 30, 60, and 120 min before fixation with 4% formaldehyde for 20 min. Before membrane permeabilization, slides were stained with mouse anti-TgSAG1 monoclonal antibody (1:1,000) for 1 hr to label attached parasites. After treatment with 0.1% Triton X-100 for 10 min, the parasites were stained with rabbit polyclonal anti-TgMIC5 antibody (1:1,000) for 1 hr to stain both invaded and attached parasites. Subsequently, slides were stained with goat anti-mouse IgG conjugated with Alexa 594 (red) (Invitrogen, 1:1,000) and goat anti-rabbit IgG conjugated with Alexa 588 (green) (Invitrogen, 1:1,000) along with DAPI for nuclear staining. After staining, slides were mounted with anti-fade Mowiol solution and observed by immunofluorescence. Extracellular parasites only showed red fluorescence, whereas intracellular parasites exhibited both red and green fluorescence. Six fields of view from individual invasion experiments were captured by a Leica^®^ DMi8 inverted epifluorescence microscope and processed with ImageJ software. The attachment efficiency of each strain was measured by dividing the total number of parasites labeled in red by the total number of host nuclei, and normalized against that of WT parasites. The invasion efficiency of each strain was quantified using the following equation ([sum of green parasites] - [sum red parasites]) / total host nuclei, and data were normalized against data for WT parasites.

### Immunofluorescence-based replication assay

Freshly egressed parasites were filter-purified and inoculated into individual wells of an 8-well chamber slide pre-seeded with HFF cells at approximately 1 × 10^5^ cells per well. Non-invaded parasites were washed off at 4 hrs post-infection. Invaded parasites were allowed to infect host cells for an additional 24 and 32 hrs before fixation. The infected host cells were stained with monoclonal anti-TgGRA7 (1:1,000) antibody and DAPI to help distinguish individual parasitophorous vacuoles (PVs) and the nuclei of parasites, respectively. Slides were subjected to standard immunofluorescence microscopy for imaging. 100 parasitophorous vacuoles were enumerated for each strain and plotted as the distribution of different sized PVs. In addition, replication was also expressed as the average number of parasites per PV.

### Luminescence-based growth assay

Parasites expressing NanoLuc luciferase were inoculated into a white 96-well tissue culture plate with a flat, solid bottom (Greiner Bio-One) pre-seeded with confluent HFF cells at 1.5 × 10^3^ cells/well. Each strain was inoculated into 4 individual wells to monitor the fold-change of luciferase activity versus time, which is proportional to intracellular growth. At 4 hrs post-infection, the individual wells were aspirated to remove non-invaded parasites. The first well was treated with 100 μL of lysis buffer containing NanoLuc luciferase substrate and incubated for 10 min, and a luminescence reading was taken by using the BioTek^®^ multimode H1 hybrid plate reader. The remainder of the 3 wells were replenished with fresh D10 medium without phenol red for an additional 24, 48, and 72 hrs. Subsequent luminescence readings were all performed via the methods above. Luminescence readings versus time were normalized against the reading at 4 hrs post-infection to calculate the fold-change of parasite growth.

### Egress assay

A Lactate dehydrogenase release assay was used to measure the egress efficiency of parasites. Freshly lysed parasites were filter-purified and resuspended in D10 medium at 5 × 10^5^ parasites/ml. 100 μL of parasite suspension was inoculated into each well of a 96-well plate pre-seeded with HFF cells. The parasites were allowed to replicate for 18-24 hrs, washed, and incubated with 50 μl of Ringer’s buffer (10 mM HEPES, pH 7.2, 3 mM HaH_2_PO_4_, 1 mM MgCl_2_, 2 mM CaCl_2_, 3 mM KCl, 115 mM NaCl, 10 mM glucose, and 1% FBS) for 20 min. Subsequently, an equal volume of 1 mM Zaprinast dissolved in Ringer’s buffer was added to the wells and incubated for 5 min at 37°C and 5% CO_2_. Uninfected wells were treated with 50 μl of Ringer’s buffer containing 1% Triton X-100 or normal Ringer’s buffer, serving as positive and negative controls, respectively. The released lactate dehydrogenase was centrifuged at 1,000 *x g* for 5 min twice to pellet insoluble cell debris. Fifty microliters of supernatant was subjected to the standard lactate dehydrogenase release assay as described previously [48]. The egress efficiency of each strain was calculated using the following equation, ([LDH activity derived from individual parasites]-[LDH activity of negative control]) / ([LDH activity of positive control]-[LDH activity of negative control]), and normalized against data for WT parasites.

### Size measurement of the VAC

The size of the VAC was quantified based on TgCPL staining (TgCPL is a VAC luminal protease). Freshly purified parasites were inoculated into pre-seeded HFF chamber slides, allowed to invade host cells for 30 min prior to fixation, stained with polyclonal rabbit anti-TgCPL antibody (1:100), and VAC diameter measured by immunofluorescence microscopy. The distance of the widest diagonal of TgCPL staining was used as the diameter of the VAC and was quantified using Leica^®^ LAS X software. Measurements for 50 individual parasites were performed for each replicate, data are presented as average ± S.D.

### Transcriptome sequencing and quantitative PCR (qPCR) assay

Total RNA was extracted from freshly lysed parasites using the Zymo^®^ Direct-zol™ RNA MiniPrep Plus kit, and converted to sequencing read libraries using the TruSeq Stranded mRNA sequencing kit (Illumina). The prepared libraries were subjected to 2 x 125 bp paired-end Illumina^®^ HiSeq2500 sequencing. Each sample was sequenced to a depth of at least 20 million reads. Differential expression profiling was performed by the Clemson University Genomics Computational Lab.

Approximately 500 ng of total RNA was used to measure the steady levels of transcripts for individual genes by using the Luna^®^ Universal One-Step RT-PCR kit (NEB). The qPCR assay was performed using the BioRad CFX96 Touch™ Real-Time PCR detection system. The quantification cycle (Cq) values for individual genes were used for double delta Cq analysis to calculate their relative abundances to that of WT parasites using the Bio-Rad^®^ CFX Maestro™ software. TgActin was used as the housekeeping gene for normalization.

### Mouse studies

Six-to eight-week-old, outbred CD-1 mice were infected by subcutaneous or intravenous injection with 100 WT or mutant parasites diluted in PBS. The infected mice were monitored for symptoms daily for a total of 30 days. Mice that appeared moribund were humanely euthanized via CO_2_ overdose, in compliance with IACUC’s approved protocol. The seroconversion of the surviving mice was tested by enzyme-linked immunosorbent assay (ELISA). The surviving mice were allowed to rest for 10 days, prior to subcutaneous injection with a challenge dose of 1000 WT parasites, and were monitored daily for survival for 30 days.

### Generation of TgCRT expression construct in yeast

*TgCRT* cDNA was PCR amplified from pTub-TgCRT-mCherry-3xmyc plasmid using a forward primer that introduced a 5’ KpnI site and *S. cerevisiae* Kozak sequence, and a reverse primer that omitted the mCherry-3xmyc tag and introduced a 3’ XmaJI site. The PCR amplified DNA was digested with KpnI and XmaJI and subcloned into pYES2-6xHis-BAD-V5 (hexa His, biotin acceptor domain, V5 tags) plasmid behind the GAL1 promoter and in front of the His-BAD-V5 epitope tags to generate the plasmid pYES/TgCRT-hbv. To generate the plasmid pYES/PMA-TgCRT-hbv, DNA encoding TgCRT-hbv was PCR amplified using a forward primer that omitted the first 900 bases of TgCRT and introduced a 5’ SacI site, and a reverse primer that included a 3’ NotI site and His-BAD-V5 tags. The amplified DNA was digested with Sacl and Notl and subcloned into a Sacl/Notl-digested pYES/PfHB3PMA (from [32]; modified via site-directed mutagenesis to introduce a SacI site at the PMA-PfCRT interface). Mutagenesis reactions were performed using reagents obtained from Agilent (Santa Clara, CA).

### Preparation of Yeast Membrane and Western blotting

Isolation of yeast membranes and detection of proteins by Western blot were as described in [20].

### Measurement of CQ incorporation in yeast expressing CRT

Quantitative growth rate analysis was used to calculate CQ transport as previously described in detail elsewhere [20,32,33]. Briefly, growth under each condition was measured in duplicate at an initial cell density of OD_600_= 0.1 in 96-well plates placed in a Tecan (Durham, NC) M200Pro or BioTek (Winooski, VT) Epoch2 plate reader. CQ-induced growth delays at 80 mM CQ, pH 6.75 were calculated as the difference in time taken to reach maximal growth rate in PMA-TgCRT non-inducing versus inducing media (see [33]).

### Statistics

Statistical analysis was performed using Prism software (GraphPad). The methods used in different assays were indicated in the figure legends.

**Table 1.** Strains used in this study.

| Name | Genetic background | Comments |
| --- | --- | --- |
| WT | RH$\Delta ku80\Delta hxg$ | Requested from the Carruthers Lab, not generated in this study |
| $\Delta crt$ | RH$\Delta ku80\Delta hxg\Delta crt$ | *TgCRT* was deleted by double crossover homologous recombination |
| $\Delta crtCRT$ | RH$\Delta ku80\Delta hxg\Delta crt::TgCRT$-<br>*mCherry-3xmyc* | Ectopic expression of a C-terminally epitope-tagged *TgCRT* in $\Delta crt$ for complementation |
| $\Delta crtCRT^{T369K}$ | RH$\Delta ku80\Delta hxg\Delta crt::TgCRT^{T369K}$-<br>*mCherry-3xmyc* | Ectopic expression of a C-terminally epitope-tagged *TgCRT* mutant in $\Delta crt$ for complementation. The original threonine at position 369 within *TgCRT* was changed to lysine by site-directed mutagenesis. |
| $\Delta crt\Delta cpb$ | RH$\Delta ku80\Delta hxg\Delta crt\Delta cpb$ | The entire *TgCPB* gene was ablated by CRISPR-Cas9 based genome editing technique |
| WT::*nLuc* | RH$\Delta ku80\Delta hxg::nLuc$ | Expressed NanoLuc luciferase in WT parasites |
| $\Delta crt::nLuc$ | RH$\Delta ku80\Delta hxg\Delta crt::nLuc$ | Expressed NanoLuc luciferase in $\Delta crt$ parasites |
| $\Delta crtCRT::nLuc$ | RH$\Delta ku80\Delta hxg\Delta crt::TgCRT$-<br>*mCherry-3xmyc::nLuc* | Expressed NanoLuc luciferase in $\Delta crtCRT$ parasites |
| TgAMN-3xHA | RH$\Delta ku80\Delta hxgTgAMN$-3xHA | *TgAMN* gene was endogenously tagged with a 3xHA epitope at its 3'-end by CRISPR-Cas9 based genome editing technique |
| TgSCP-3xmyc | RH$\Delta ku80\Delta hxgTgSCP$-3xmyc | *TgSCP* gene was endogenously tagged with a 3xmyc epitope at its 3'-end by single crossover recombination |

## Data Availability Statement

The raw data of transcriptomic sequencing of WT and Δ*crt* strains have been deposited in the NIH Gene Expression Omnibus (GEO) database. The accession number is GSE116539. https://www.ncbi.nlm.nih.gov/geo/query/acc.cgi?acc=GSE116539

## Acknowledgments

We thank our colleagues Drs. Michael Blackman, Peter Bradley, Vern Carruthers, Silvia Moreno, and David Sibley for kindly providing key reagents for this study. We also want to thank Drs. Rooksana Noorai and Vijay Shankar at Clemson University Genomics Computational Lab for providing technical assistance and expertise in performing transcriptomic sequencing and differential gene expression analysis and depositing the raw data files of transcriptomic sequencing into the NIH GEO database. We acknowledge Dr. Lesly Temesvari for critically reading this manuscript before submission.

## Author Contributions

**Conceptualization**: Paul D. Roepe, Zhicheng Dou.

**Formal analysis**: L. Brock Thornton, Paige Teehan, Katherine Floyd, Christian Cochrane, Amy Bergmann, Bryce Riegel, Paul D. Roepe, Zhicheng Dou.

**Funding acquisition**: Paul D. Roepe, Zhicheng Dou.

**Investigation**: L. Brock Thornton, Paige Teehan, Katherine Floyd, Christian Cochrane, Amy Bergmann, Bryce Riegel, Zhicheng Dou.

**Methodology**: L. Brock Thornton, Paige Teehan, Bryce Riegel, Paul D. Roepe, Zhicheng Dou.

**Project administration**: Paul D. Roepe, Zhicheng Dou.

**Resources**: Paige Teehan, Amy Bergmann, Bryce Riegel, Zhicheng Dou.

**Supervision**: Paul D. Roepe, Zhicheng Dou.

**Writing - original draft**: Zhicheng Dou.

**Writing - review & editing**: L. Brock Thornton, Bryce Riegel, Paul D. Roepe, Zhicheng Dou.

---

**Figure S1. No defects in intramembrane proteolytic cleavage of micronemal protein were observed in the Δ*crt* mutant**. Purified, extracellular parasites that had not been permeabilized were stained with anti-TgMIC2 and anti-TgSAG1 antibodies in order to measure the retention of TgMIC2 on the parasite surface. During secretion, the TgMIC2 protein is cleaved by intramembrane rhomboid proteases, such as TgROM4. The abundance of TgMIC2 on the surface of Δ*crt* parasites was similar to that of the WT and Δ*crtCRT* strains, indicating that there is comparable intramembrane cleavage of TgMIC2 among the parasites with or without TgCRT.

**Figure S2. Schematic of the endogenous epitope-tagging of *Toxoplasma* putative aminopeptidase N (TgAMN, TGGT1_221310) and putative Pro-Xaa serine carboxypeptidase (TgSCP, TGGT1_254010)**. **A)** The plasmid encoding Cas9 and sgRNA targeting TgAMN was co-transfected into WT parasites with the PCR product carrying a 3xHA epitope tag and a pyrimethamine resistance cassette (DHFR) flanked by 50 bp regions upstream and downstream of the stop codon of TgAMN. The 3xHA tag and the drug resistance cassette were incorporated at the C-terminus of the *Toxoplasma* putative aminopeptidase N via double crossover homologous recombination mediated by the CRISPR-Cas9 genome editing tool. **B)** The putative Pro-Xaa serine carboxypeptidase was endogenously tagged with a 3xmyc epitope tag at its C-terminus by single crossover homologous recombination. A 1 kb region upstream of the stop codon of TgSCP was amplified and fused at the 5’-end of the 3xmyc tag to produce the TgSCP-3xmyc tagged plasmid. The 1 kb TgSCP-coding region was cleaved by an endonuclease in the middle prior to transfection to facilitate its integration.

**Figure S3. The subcellular concave structure in the Δ*crt*Δ*cpb* mutant was shrunken relative to the Δ*crt* strain during the extracellular stage**. WT, Δ*crt, ΔcrtCRT*, and Δ*crt*Δ*cpb* parasites were purified and attached on the surface of a slide for differential interference contrast (DIC) microscopy imaging. Although the Δ*crt*Δ*cpb* mutant still showed an enlarged concave subcellular structure (indicated by the arrow), their sizes were significantly smaller than those in the Δ*crt* mutant. Scale bar = 5 μm.

**Figure S4. Alignment of TgCRT and PfCRT primary sequences and schematic of the PMA-TgCRT construct expressed in *S. cerevisiae*. A)** Alignment of TgCRT and PfCRT amino acid sequences reveals the 300 most N-terminal residues to be non-homologous, and that they do not encode any putative transmembraneous domains or inter helical loop regions, whereas the remainder of TgCRT is highly homologous to PfCRT. Alignment analysis also revealed that the threonine residue at position 369 within TgCRT corresponds to the well-characterized lysine residue at position 76 within PfCRT (highlighted in red box). Identical and similar residues are highlighted in black and dark grey, respectively. **B)** The 111 most N-terminal residues of *S. cerevisiae* plasma membrane ATPase (PMA; black) are fused in frame to the truncated TgCRT (dark grey) from which the first 300 codons have been deleted. The construct includes a C-terminal tag comprised of hexaHIS (H), biotin acceptor domain (B), and V5 epitope tag (V; “HBV” light grey). **C)** Primary amino acid structure of PMA-TgCRT. Residues from PMA are shown in black, those from TgCRT are in shown dark grey, and those comprising the tag are shown in light grey.

**Figure S5. Anti-V5 Western blot analysis of TgCRT and PfCRT constructs expressed in *S. cerevisiae***. Each lane contains 40 μg of protein. Lane 1, yeast membranes for yeast expressing empty vector (EV); lane 2, PfCRT membranes; lane 3, TgCRT membranes; lane 4, PMA-TgCRT fusion membranes; lane 5, blank; lane 6, cytosol from TgCRT yeast; lane 7, cytosol from PMA-TgCRT yeast. The unmodified TgCRT is not expressed in *S. cerevisiae* (lane 3), however, the PMA-TgCRT fusion construct is expressed to similar levels relative to PfCRT [20], and is membrane localized. Lower molecular mass bands in lane 4 are proteolytic products, and can also be found in the cytosolic fraction (lane 7).

**Figure S6. Δ*crt* parasites were more sensitive to the treatment of chloroquine than WT parasites**. WT, Δ*crt*, and Δ*crtCRT* parasites expressing luciferase were used to infect host cells in the presence of 100 μM or 0 μM chloroquine. The luciferase activity of each strain was measured at 2 and 26 hrs postinfection. The ratios of the luciferase activities determined at 26 hours over that at 2 hrs were plotted. Statistical significance was determined using unpaired Student’s t-test. *, *p*<0.05; n.s., not significant.

**Table S1. Differential gene expression analysis between WT and Δ*crt* strains**. The genes whose fold changes were >1.5 and p-values were <0.05 were listed.

**Table S2. Primers used in this study**.

